# Tracking chromatin state changes using *μ*Map photo-proximity labeling

**DOI:** 10.1101/2021.09.28.462236

**Authors:** Ciaran P. Seath, Antony J. Burton, David W. C. MacMillan, Tom W. Muir

## Abstract

Interactions between biomolecules, particularly proteins, underlie all cellular processes, and ultimately control cell fate. Perturbation of native interactions through mutation, changes in expression levels, or external stimuli leads to altered cellular physiology and can result in either disease or therapeutic effects.^1,2^ Mapping these interactions and determining how they respond to stimulus is the genesis of many drug development efforts, leading to new therapeutic targets and improvements in human health.^1^ However, in the complex environment of the nucleus it is challenging to determine protein-protein interactions due to low abundance, transient or multi-valent binding, and a lack of technologies that are able to interrogate these interactions without disrupting the protein binding surface under study.^3^ Chromatin remodelers, modifying enzymes, interactors, and transcription factors can all be redirected by subtle changes to the microenvironment, causing global changes in protein expression levels and subsequent physiology. Here, we describe the Chroma-*μ*Map method for the traceless incorporation of Ir-photosensitizers into the nuclear microenvironment using engineered split inteins. These Ir-catalysts can activate diazirine warheads to form reactive carbenes within a ~10 nm radius, cross-linking with proteins within the immediate microenvironment for analysis via quantitative chemoproteomics.^4^ We demonstrate this concept on nine different nuclear proteins with varied function and in each case, elucidating their microenvironments. Additionally, we show that this short-range proximity labeling method can reveal the critical changes in interactomes in the presence of cancer-associated mutations, as well as treatment with small-molecule inhibitors. Chroma-*μ*Map improves our fundamental understanding of nuclear protein-protein interactions, as well as the effects that small molecule therapeutics have on the local chromatin environment, and in doing so is expected to have a significant impact on the field of epigenetic drug discovery in both academia and industry.

## Main Text

Mapping protein-protein interactions (PPIs) is central to our understanding of cellular biology.^5^ The enormous challenges associated with this undertaking are magnified in the nucleus where transient and multivalent interactions, fine-tuned by post-translational modifications (PTMs), combine to choreograph DNA-templated processes such as transcription.^6^ Perturbations to these regulatory mechanisms often leads to disease,^7^ for example, somatic mutations that alter the composition and activity of chromatin-associated protein complexes are implicated in many human cancers and developmental disorders.^8 9,10^ Moreover, recent studies have shown that histone proteins are themselves frequently mutated in cancers.^11–13^ Understanding how these mutations lead to, or perpetuate disease, is the focus of intense investigation,^11–13^ work that necessitates accurate comparative mapping of chromatin-associated PPI networks as a function of altered cell states.^1^

The elucidation of nuclear PPIs has typically been performed by immunoprecipitation/mass spectrometry (IP/MS) workflows, where antibodies recognizing selected proteins are used to enrich their target along with direct interactors.^14^ However, IP/MS approaches rely on nuclear lysates as input, which may not be ideal for every system, ^15,16^ especially when the interactions are transient in nature (e.g. PTM-driven) or require multi-protein complexes that bridge native chromatin.^17,18,19^ This has fuelled the development of chemo-proteomics approaches such as those based on photocrosslinking^20,21^ or proximity labeling^22–24^ technologies that seek to capture PPIs in a native-like environment. Despite these ongoing advances, no single method exists to map chromatin interactomes in a general and unbiased manner, and particularly to discern how such interactions are affected by perturbations such as mutation or drug treatment (Figure 1a). To address this need we envisioned the union of technologies recently disclosed by our two laboratories, *μ*Map^4^ and *in nucleo* protein *trans*-splicing,^20^ to enable a traceless, short-range proximity labeling method that can be readily deployed to any nuclear protein target. Protein *trans*-splicing using ultra-fast and bioorthogonal split inteins facilitates the installation of Ir-photocatalysts onto the N- or C-termini of target proteins. Upon irradiation with blue LEDs in the presence of a biotin-diazirine probe, localized carbene generation allows interactomes to be determined, specifically those within ~10 nm of the iridium-centered photocatalyst (Figure 1b).

**Figure 1.**
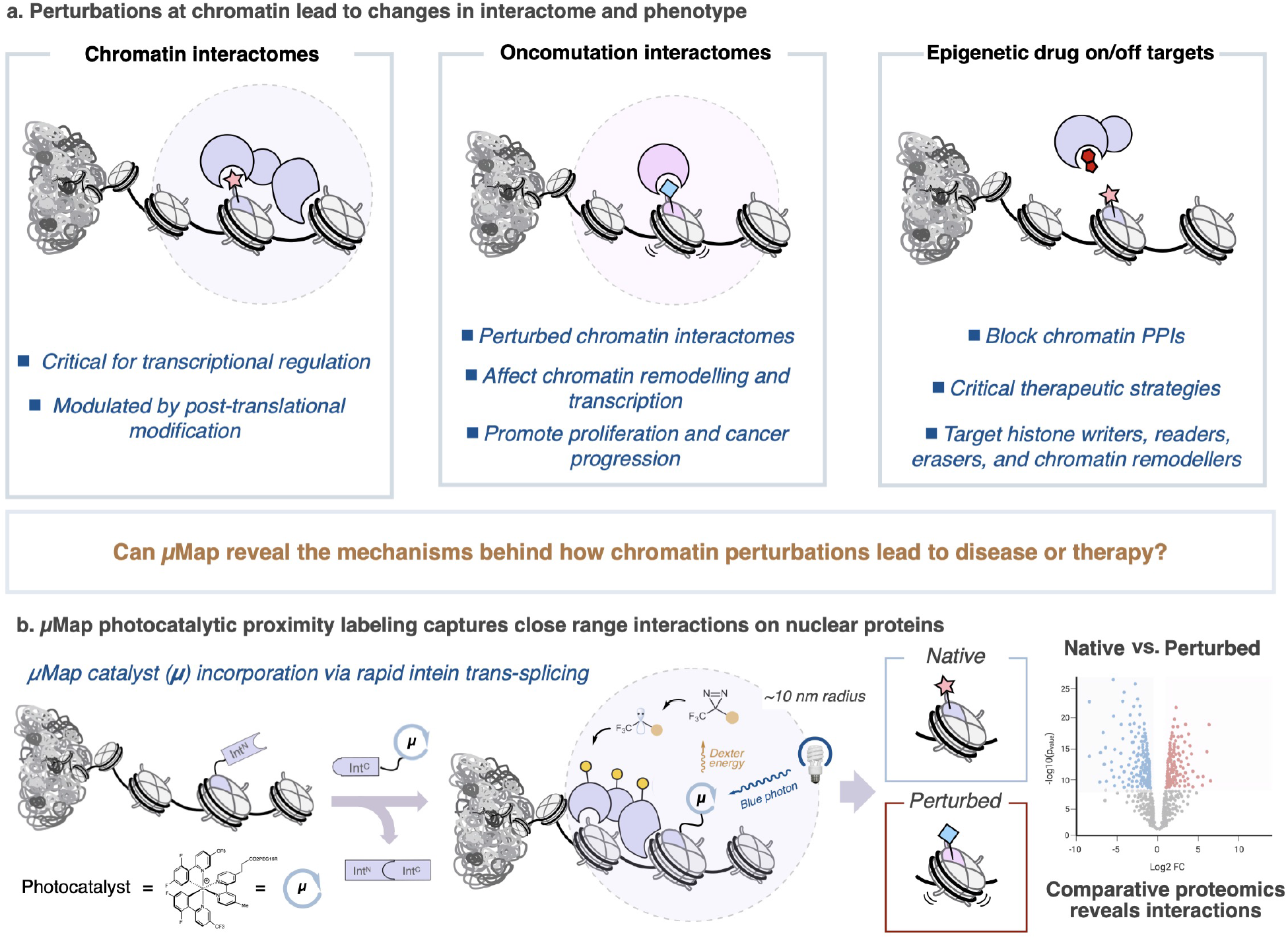
Development of a catalytic labeling platform using inteins. **a**, Chromatin interactomes can be perturbed through mutation leading to oncogenic phenotypes. Epigenetic drugs alter the chromatin interactome for therapeutic benefit. **b,** Cartoon showing strategy for nuclear photoproximity labeling. Ir-photocatalysts can be incorporated into nuclear proteins via protein *trans-splicing*. The N-terminal fragment of the engineered Cfa split intein (Cfa^N^) is fused to a target nuclear protein, while the complementary C-terminal fragment (Cfa^C^) is linked to the photocatalyst. *In nucleo* splicing provides Ir-conjugated nuclear proteins. Comparative chemoproteomic analysis reveals how a given perturbation (e.g. a mutation or drug treatment) affects local interactomes, providing insights into the mechanistic basis for disease or therapy.

Our proposed workflow offers distinct advantages for the elucidation of subtly perturbed chromatin interactomes through mutation or ligand binding. Principally, the short radius afforded by this technology limits labeling to the close vicinity of a designated chromatin factor or nucleosome, only identifying proteins that are affected by a particular mutation or pharmacological intervention. This is critically important given the structural and functional heterogeneity of chromatin.^25^ Furthermore, the incorporation of the *μ*Map catalyst is designed to be almost traceless and its small size is expected to minimize disruption to the native environment (e.g. in comparison to fusion proteins), allowing the study of modifications that only minimally change the native interactome.

We began our studies by fusing the N-terminal fragment of the engineered Cfa split intein (Cfa^N^)^26,27^ to the C-terminus of histone H3.1. For analytical convenience, we also included HA and FLAG epitope tags flanking the intein (Figure 2a). Expression of this construct (H3.1-HA-Cfa^N^-FLAG) in HEK293T cells led to its incorporation into chromatin (Figure 2b). We then treated nuclei isolated from these transfected cells with the complementary split intein fragment (Cfa^C^) linked to the iridium photocatalyst (Figure S1). This resulted in the site-selective incorporation of the photocatalyst onto the C-terminus of H3.1 through *in nucleo* protein trans-splicing (Figure 2b). Irradiation of these nuclei in the presence of the diazirine-biotin probe (Dz-Bt) led to dramatically enhanced protein labeling compared to a control reaction in which H3.1-HA-Cfa^N^-FLAG expressing nuclei were treated with a free Ir-photocatalyst (Figure 2b). Importantly, elution from streptavidin beads showed strong enrichment of histone H3 by western blot (Figure 2b).

**Figure 2.**
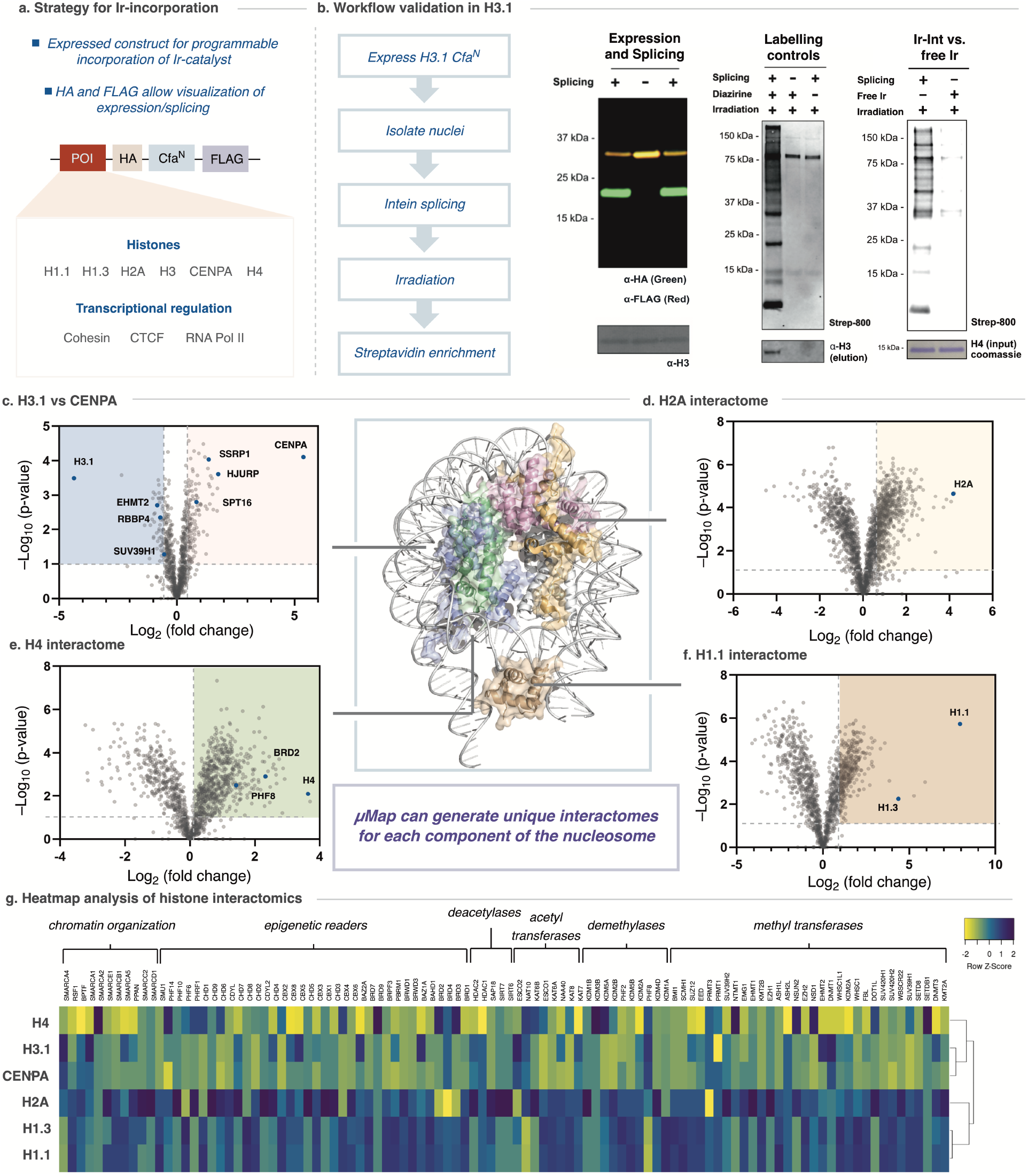
Chroma-μMap proximity labeling can discriminate between histone interactomes. **a,** Programmable incorporation of iridium photocatalysts into proteins of interest (POI) can be achieved by transfection of POI-HA-Cfa^N^-FLAG constructs. **b,** Validation of the approach using histone H3. *In nucleo* protein *trans*-splicing with 0.5 μM Cfa^C^-Ir for 1 h results in the generation of the H3 spliced product (green) as shown by western blotting (left panel). Biotinylation of nuclear proteins, as detected by streptavidin blotting, is dependent on irradiation, Cfa^C^-Ir and the diazirine probe (middle, right panels). H3 is enriched following labeling and streptavidin enrichment. **c-f,** Volcano plots displaying proteins enriched in a series of comparative proteomics studies using the Chroma-μMap workflow. Select proteins discussed in the main text are highlighted. Panel c shows proteins enriched in H3.1-Ir vs CENPA-Ir, whereas in the other panels the comparison is made with the free iridium catalysts added to the cells (i.e. not conjugated to the POI). The structure of the nucleosome (pdb code: 7K5X) in the center is color-coded to match the histone proteins under investigation. FDR values were calculated using the Benjamini–Hochberg procedure, as described in the Methods. **g,** Heatmap analysis of chromatin-associated protein enrichment across the 6 histones studied vs. free Ir-catalyst.

Encouraged by this, we established a tandem mass tag (TMT)-based quantitative chemo-proteomics workflow to determine the interactome for H3.1 versus a free Ir control and the centromere-specific H3 variant CENP-A (Figure 2c, Figure S2&3 & Table S1–3). Each Chroma-*μ*Map experiment was performed in triplicate, with the Ir-appended protein of interest enriched in each case. Comparison of the H3.1 vs CENP-A interactomes returned established H3.1 modifying enzymes (e.g. EHMT2, SUV39H1) and reader proteins (e.g. HP1 isoforms) (Figure 2c,g, Table S1–3).^28^ CENP-A interacting proteins included HJURP and both members of the FACT complex (SSRP1 & SPT16), responsible for the deposition of CENP-A into chromatin (Figure 2c).^29,30^ Interactomes for histone H4 and H2A were also obtained versus the free Ir control (Figure 2d, e, g, Figure S4&5 and Table S4&5), with histone-specific interactors observed (e.g. BRD2 & PHF8 for H4).

We then extended our analysis to the linker histone H1. Traditionally considered a chromatin architectural protein, there is emerging evidence that this nucleosome binding protein can also function as a PPI hub, although this remains an understudied area of chromatin biology.^31^ Here, we compared the interactomes of two linker histone isoforms, H1.1 and H1.3, along with the free Ir control (Figure 2f,g, Figure S6–9, Table S6&7). We found 506 shared hits between the two isoforms, consistent with the high sequence homology between the proteins. Fewer unique interactors were observed, with 33 and 27 unique proteins observed for H1.1 and H1.3, respectively (Figure S8, Table S6&7). Heatmap analysis of these data showed significant differences across all six histones studied (Figure 2g, Figure S10), suggesting that *μ*Map can provide interactome data for distinct protein surfaces within a short radius. For example, we observed that histone H2A clustered more closely with the linker histones than to H4 or the H3 variants, perhaps reflecting the position of the H2A C-terminus (i.e. where the Ir photocatalyst is attached) at the DNA entry-exit site, the region where the globular domain of H1 also engages the nucleosome^32^ (Figure S11).

Next, we expanded the scope of the Chroma-*μ*Map workflow to include the nuclear proteins SMC1A, a member of the cohesin complex, and the transcription factor CTCF (Figure S12–15, Table S8&9). These photo-proximity labeling analyses revealed enrichment of the bait proteins in addition to known interactors (e.g. SMC1B and WAPAL for SMC1A, and MAFb for CTCF)^33,34^. Intriguingly, BHLHE40,^35,36^ a basic helix-loop-helix transcription factor implicated in circadian rhythm^37^ and immunity^38^, was highly enriched in both datasets, suggesting it may play a role in cohesin/CTCF based chromatin reorganisation.^35^

With the basic chemo-proteomics workflow established, we next attempted to map how histone PPIs are perturbed through mutation. Recent sequencing of patient tumour samples identified >4,000 histone mutations associated with a wide range of cancers.^11–13,39^ Determining whether such mutations drive oncogenesis and cancer progression, or are simply passenger mutations, is critical to identifying novel opportunities for therapeutic intervention. We sought to apply our method to the cancer-associated histone mutation H2A E92K, which is correlated with oesophageal and stomach cancers and aggressive disease progression.^13^ This mutation introduces a charge swap within the critical acidic patch interaction motif on the nucleosome, into which arginine residues of interacting proteins are known to anchor.^40–43^ We hoped that our methodology could shed light on to what extent chromatin PPIs are perturbed by this oncohistone mutation (Figure 3a).

**Figure 3.**
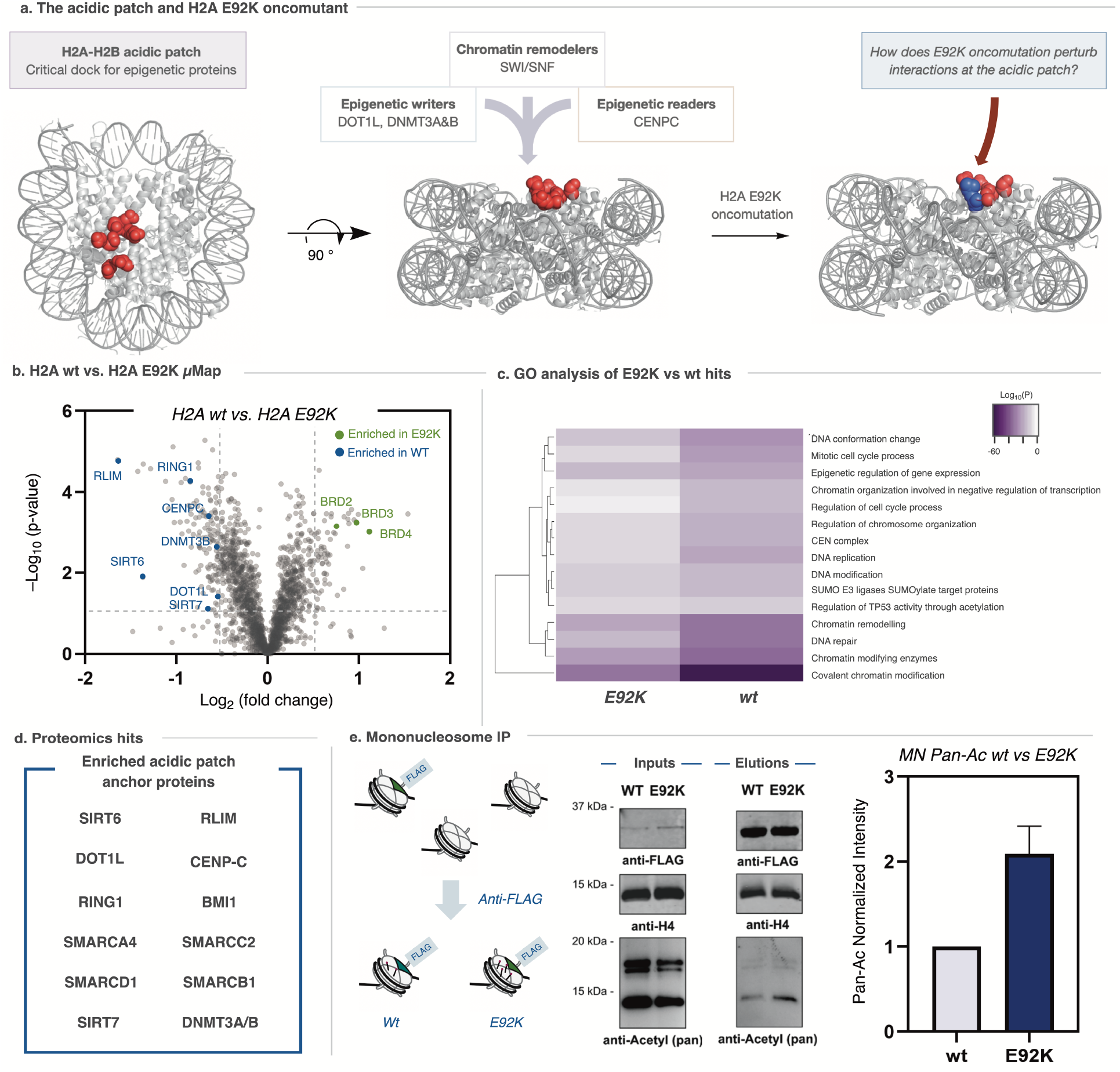
μMap as a method to uncover oncogenic function of the somatic mutation H2A E92K. **a,** The histone acidic patch provides a crucial dock for chromatin regulatory complexes. Such complexes exploit an arginine “anchor” residue to anchor to the nucleosome surface (pdb: 1kx5). H2A E92K is associated with stomach and oesophageal cancers and alters the electrostatic potential of the acidic patch. **b,** A volcano plot displaying protein interactors from the Chroma-μMap method comparing H2A vs H2A E92K. Select proteins are highlighted. FDR values were calculated using the Benjamini–Hochberg procedure, as described in the Methods. **c,** Comparative GO analysis for wt vs E92K hits. Consistent with the role of the acidic patch, GO terms are deenriched in the E92K mutant. **d,** Selected proteins enriched by wt H2A over H2A E92, consistent with E92K disrupting binding of chromatin affectors to the acidic patch. **e,** Increased local acetylation of histone H4 is observed in the presence of H2A E92K, as determined by mononucleosome immunoprecipitation experiments. Western blot analysis showed an increase (2.1 +/− 0.33 error) in acetylation in the E92K mutant compared to wild type. Histogram shows change in acetylation levels on H4 normalized to wild-type as determined by densitometry analysis of western blots (s.e.m. n=4).

We found the local chromatin microenvironment is indeed sensitive to the H2A E92K mutation (Figure 3b,d, Figure S4, Table S10). GO analysis revealed diminished enrichment of proteins related to chromatin modification and reorganization as a function of the perturbed acidic patch (Figure 3c). Intriguingly, SIRT6, a deacetylase, was enriched in the wild-type sample, whereas the transcriptional activators, BRD2/3/4 were all enriched by E92K. Taken together, this suggests the mutation acts to block histone lysine deacetylase activity, resulting in increased local acetylation and binding of associated reader proteins. To support this hypothesis, we expressed both wild-type and H2A E92K histones in HEK 293T cells and isolated intact mononucleosomes by anti-FLAG immunoprecipitation (Figure 3e). We found global lysine acetylation was increased 2-fold in the mutant nucleosomes over wild-type, with a particular increase observed on H4. This effect is striking given that we are likely only enriching one copy of the mutant histone per nucleosome due to presumed stochastic incorporation.^44,45^ These data demonstrate that the resolution provided by our proximity labeling method can be used to uncover molecular level details for gain-of-function or loss-of-function interactions in the nucleus in a single experiment.

Excited by this, we posited that our method could be used to determine the roles of small molecule ligands in the chromatin microenvironment (Figure 4a). Epigenetic drug discovery has become a critical focus for therapeutic intervention, encompassing dozens of targets across a range of therapeutic areas. ^3^ We began by examining the effects that the bromodomain inhibitor JQ-1 has on chromatin (Figure 4b). JQ-1 is known to target the BET family of bromodomain containing readers (BRD2/3/4), with analogues progressing through the clinic.^46,47^ We theorized that Chroma-*μ*Map would be sensitive enough to measure the effects that BRD inhibition would have on the chromosomal microenvironment. To probe this, we compared the interactomes of Ir-conjugated H2A in the presence or absence of 5 μM JQ-1 (Figure 4b, Figure S16, Table S11). BRD2/3/4 were all significantly enriched in the untreated sample, consistent with blocking bromodomainnucleosome interactions. The known JQ-1 off-target SOAT1^48^ was also enriched in the untreated sample.

**Figure 4.**
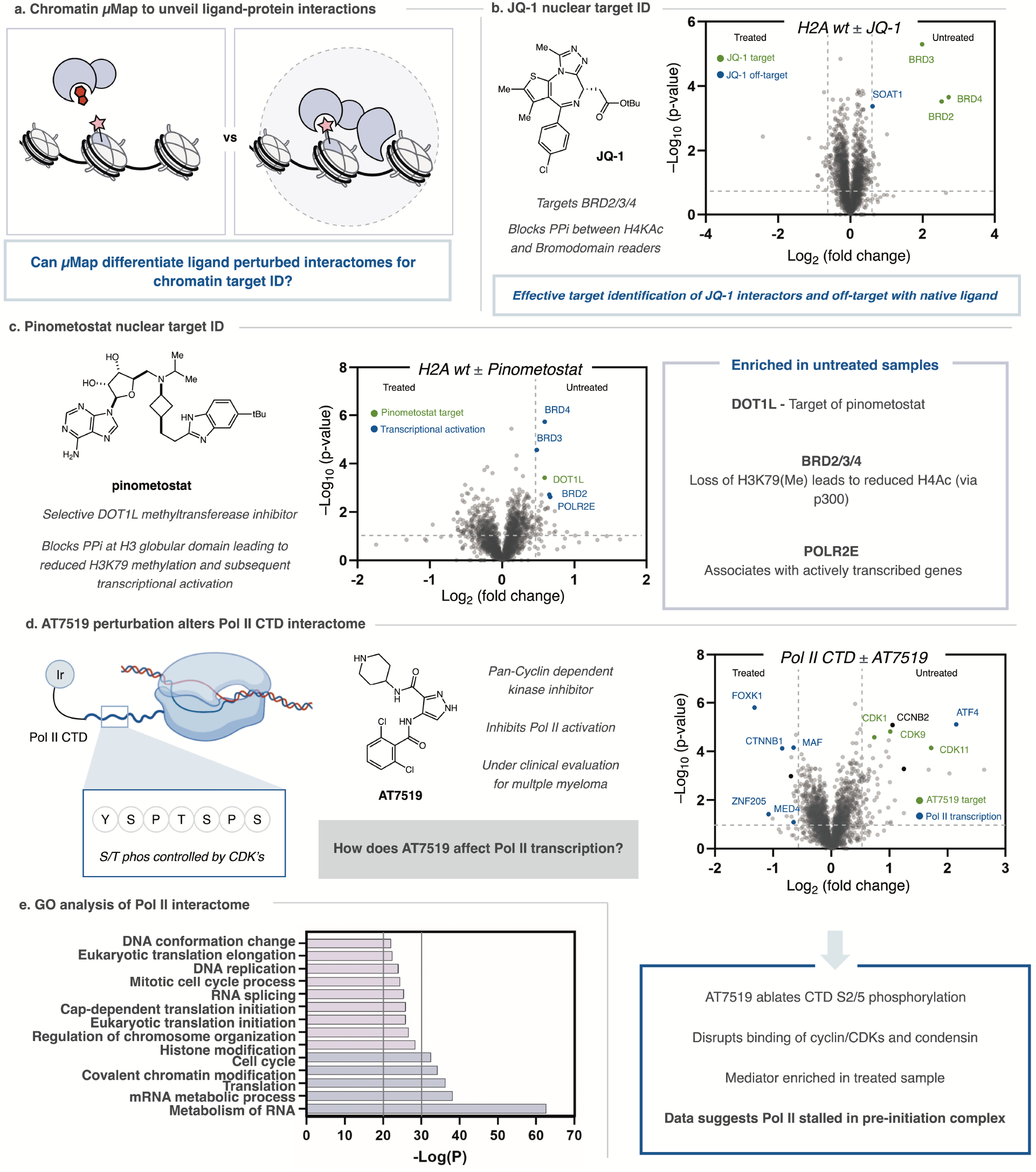
Interactome mapping of nuclear proteins to assess ligand interactions. **a,** Performing the Chroma-*μ*Map workflow in the presence of a bioactive ligand unveils protein-protein interactions that are disrupted or promoted by ligand treatment. This method can provide nuclear target identification data. **b,** Volcano plot displaying the H2A interactome vs H2A + 10 μM JQ-1 (structure shown at left). This method identifies BRD2/3/4 as JQ-1 target proteins in addition to known off-target SOAT1. **c,** Volcano plot displaying the H2A interactome vs H2A + 2.5 μM pinometostat. Pinometostat target DOT1L is enriched, in addition to several proteins associated with loss of H3K79me. **d,** RNA Pol II transcription is controlled by phosphorylation of the CTD. AT7519 inhibits CTD phosphorylation, stalling transcription. Volcano plot displaying RPB1-Ir vs RPB1-Ir + 5 μM AT7519. FDR values were calculated using the Benjamini–Hochberg procedure, as described in the Methods. **e,** GO analysis of POL II interactome indicates the identified interactome is significantly related to the annotated role of RNA Pol II. Analysis performed on hits from RPB1-Ir vs Ir.

We then performed a similar experiment with the DOT1L methyltransferase inhibitor, pinometostat, to assess both the selectivity of this ligand and the effect that depletion of H3K79 methylation has on the chromatin microenvironment.^49,50^ DOT1L was enriched in the untreated sample, as were several proteins related to transcriptional activation, including BRD2/3/4 and POLR2E (Figure 4c. Figures S16&17, Table S12). This observation is consistent with previous reports demonstrating that H3K79 methylation leads to recruitment of the acetyltransferase P300, subsequent BRD recruitment, and transcriptional activation.^50^ It was gratifying to see that in a single experiment we can extract both target-ID data for a small molecule ligand along with downstream transcriptional effects.

Finally, we applied our method to RNA polymerase II (POL II), which is responsible for transcribing protein-encoding genes in a highly regulated process involving the sequential association of multiple protein complexes.^51–53^ Release of promoter-proximal paused POL II is primarily achieved through phosphorylation of the disordered C-terminal domain (CTD) of the RPB1 subunit, comprised of a repeating unit of Y-S-P-T-S-P-S, by cyclin dependent kinases (CDKs). Inhibition of CDKs has therefore become an attractive therapeutic strategy to stall POL II and downregulate transcription and therefore cancer proliferation. The small molecule ligand AT7519, a pan-CDK inhibitor, has been developed to probe this strategy in the context of multiple myeloma and is currently in phase 1 clinical trials (Figure S18).^54^ We questioned if we could determine how this ligand affects the POL II interactome, and whether our method could unveil at what stage in its transcriptional cycle POL II is arrested by AT7519 treatment.

To assess this, we expressed RPB1-Cfa^N^ in HEK 293T cells, where the intein is fused to the C-terminus of the CTD (Figure S19&22). The Chroma-*μ*Map workflow was then performed in the presence and absence of AT7519 (Figure 4d, Figure S20, Table S13). Ontology analysis of the RPB1 interactome (vs. free Ir-catalyst) showed good agreement with the known functions of POL II (Figure 4e, Figure S21, Table S14), with metabolism of RNA being highly enriched (>60 −Log P-value). Comparing our treated and untreated datasets revealed CDK11, 9, and 1 to be enriched in the untreated sample, consistent with wide-ranging CDK inhibition by AT7519 (Figure 4d).^54^ Mediator complex subunits in addition to several transcription factors were enriched in the treated sample, suggesting CDK inhibition halts the progression of POL II at the pre-initiation complex, before CTD phosphorylation and association of the NELF and DSIF complexes.^55^ Intriguingly, cyclin T1, a member of the TEF complex was also enriched in the AT7519 treated sample, despite its binding partner CDK9 showing the opposite enrichment.

## Conclusion

In summary, we have developed a photocatalytic proximity labeling technology that can be deployed across the nuclear proteome. The short range diazirine activation mechanism allows the collection of highly precise interactomics data that is sensitive to single amino acid mutations and can be used to detect changes caused by external stimuli such as ligand incubation. We believe this method will be broadly applicable across nuclear biology for the study of disease-associated mutations. Additionally, this method is an effective tool for ligand target-ID in chromatin, identifying on-and off-target proteins, and revealing how treatment with these molecules impacts local chromatin interactomes.

## Supporting information

Full MS Processed Data

**Supplementary Information** is linked to the online version of the paper.

## Data availability

All relevant data are included in the manuscript and supplementary information.

## Acknowledgements

Research reported in this publication was supported by the NIH National Institute of General Medical Sciences (R35GM134897-01, R01-GM103558-03 and R37-GM086868) and the NIH National Cancer Institute (P01 CA196539). AJB was a Damon Runyon Fellow of the Damon Runyon Cancer Research Foundation (DRG-2283-17). The authors thank Saw Kyin and Henry H. Shwe at the Princeton Proteomics Facility.

## Author Contributions

CPS, AJB, TWM, DWCM conceived the work. CPS, AJB, TWM, DWCM designed and executed the experiments. CPS, AJB, TWM, DWCM prepared this manuscript.

## Author Information

The authors declare no competing financial interests. Readers are welcome to comment on the online version of the paper.

## References

1. Scott, D. E., Bayly, A. R., Abell, C. & Skidmore, J. Small molecules, big targets: drug discovery faces the protein–protein interaction challenge. Nat. Rev. Drug Discov. 15, 533–550 (2016).

2. Veltman, J. A. & Brunner, H. G. De novo mutations in human genetic disease. Nat. Rev. Genet. 13, 565–575 (2012).

3. Campbell, R. M. & Tummino, P. J. Cancer epigenetics drug discovery and development: The challenge of hitting the mark. J. Clin. Invest. 124, 64–69 (2014).

4. Geri, J. B. et al. Microenvironment mapping via Dexter energy transfer on immune cells. Science (80-.). 367, 1091–1097 (2020).

5. Ruffner, H., Bauer, A. & Bouwmeester, T. Human protein-protein interaction networks and the value for drug discovery. Drug Discov. Today 12, 709–716 (2007).

6. Kouzarides, T. Chromatin Modifications and Their Function. Cell 128, 693–705 (2007).

7. Ganesan, A., Arimondo, P. B., Rots, M. G., Jeronimo, C. & Berdasco, M. The timeline of epigenetic drug discovery: from reality to dreams. Clin. Epigenetics 11, 174 (2019).

8. Schick, S. et al. Systematic characterization of BAF mutations provides insights into intracomplex synthetic lethalities in human cancers. Nat. Genet. 51, 1399–1410 (2019).

9. Cheng, F. et al. Comprehensive characterization of protein–protein interactions perturbed by disease mutations. Nat. Genet. 53, 342–353 (2021).

10. Cheng, Y. et al. Targeting epigenetic regulators for cancer therapy: Mechanisms and advances in clinical trials. Signal Transduct. Target. Ther. 4, (2019).

11. Weinberg, D. N., Allis, C. D. & Lu, C. Oncogenic mechanisms of histone H3 mutations. Cold Spring Harb. Perspect. Med. 7, 1–14 (2017).

12. Bagert, J. D. et al. Oncohistone mutations enhance chromatin remodeling and alter cell fates. Nat. Chem. Biol. 17, 403–411 (2021).

13. Nacev, B. A. et al. The expanding landscape of ‘oncohistone’ mutations in human cancers. Nature 567, 473–478 (2019).

14. Müller, M. M. & Muir, T. W. Histones: At the crossroads of peptide and protein chemistry. Chem. Rev. 115, 2296–2349 (2015).

15. Vermeulen, M. & Déjardin, J. Locus-specific chromatin isolation. Nat. Rev. Mol. Cell Biol. 21, 249–250 (2020).

16. Van Mierlo, G. & Vermeulen, M. Chromatin proteomics to study epigenetics - Challenges and opportunities. Mol. Cell. Proteomics 20, 100056 (2021).

17. Ciferri, C. et al. Molecular architecture of human polycomb repressive complex 2. Elife 2012, 1–22 (2012).

18. Ruthenburg, A. J. et al. Recognition of a mononucleosomal histone modification pattern by BPTF via multivalent interactions. Cell 145, 692–706 (2011).

19. Zhao, S., Yue, Y., Li, Y. & Li, H. Identification and characterization of ‘readers’ for novel histone modifications. Curr. Opin. Chem. Biol. 51, 57–65 (2019).

20. Burton, A. J. et al. In situ chromatin interactomics using a chemical bait and trap approach. Nat. Chem. 12, 520–527 (2020).

21. Kleiner, R. E., Hang, L. E., Molloy, K. R., Chait, B. T. & Kapoor, T. M. A Chemical Proteomics Approach to Reveal Direct Protein-Protein Interactions in Living Cells. Cell Chem. Biol. 25, 110–120.e3 (2018).

22. Seath, C. P., Trowbridge, A. D., Muir, T. W. & Macmillan, D. W. C. Reactive intermediates for interactome mapping. Chem. Soc. Rev. 50, 2911–2926 (2021).

23. Villaseñor, R. et al. ChromID identifies the protein interactome at chromatin marks. Nat. Biotechnol. 38, 728–736 (2020).

24. Ummethum, H. & Hamperl, S. Proximity Labeling Techniques to Study Chromatin. Front. Genet. 11, 1–13 (2020).

25. Baldi, S., Korber, P. & Becker, P. B. Beads on a string—nucleosome array arrangements and folding of the chromatin fiber. Nat. Struct. Mol. Biol. 27, 109–118 (2020).

26. Stevens, A. J. et al. A promiscuous split intein with expanded protein engineering applications. Proc. Natl. Acad. Sci. U. S. A. 114, 8538–8543 (2017).

27. Stevens, A. J. et al. Design of a Split Intein with Exceptional Protein Splicing Activity. J. Am. Chem. Soc. 138, 2162–2165 (2016).

28. Scott, W. A. & Campos, E. I. Interactions With Histone H3 & Tools to Study Them. Front. Cell Dev. Biol. 8, 1–21 (2020).

29. Pan, D. et al. Mechanism of centromere recruitment of the CENP-A chaperone HJURP and its implications for centromere licensing. Nat. Commun. 10, 1–18 (2019).

30. Chen, C. C. et al. Establishment of Centromeric Chromatin by the CENP-A Assembly Factor CAL1 Requires FACT-Mediated Transcription. Dev. Cell 34, 73–84 (2015).

31. Kalashnikova, A. A., Rogge, R. A. & Hansen, J. C. Linker histone H1 and protein-protein interactions. Biochim. Biophys. Acta - Gene Regul. Mech. 1859, 455–461 (2016).

32. Fyodorov, D. V., Zhou, B. R., Skoultchi, A. I. & Bai, Y. Emerging roles of linker histones in regulating chromatin structure and function. Nat. Rev. Mol. Cell Biol. 19, 192–206 (2018).

33. Stik, G. et al. CTCF is dispensable for immune cell transdifferentiation but facilitates an acute inflammatory response. Nat. Genet. 52, 655–661 (2020).

34. Petela, N. J. et al. Scc2 Is a Potent Activator of Cohesin’s ATPase that Promotes Loading by Binding Scc1 without Pds5. Mol. Cell 70, 1134–1148.e7 (2018).

35. Hu, G. et al. Systematic screening of CTCF binding partners identifies that BHLHE40 regulates CTCF genome-wide distribution and long-range chromatin interactions. Nucleic Acids Res. 48, 9606–9620 (2020).

36. Rauschmeier, R. et al. Cell-intrinsic functions of the transcription factor Bhlhe40 in activated B cells and T follicular helper cells restrain the germinal center reaction and prevent lymphomagenesis. bioRxiv 2021.03.12.435122 (2021) doi:10.1101/2021.03.12.435122.

37. Honma, S. et al. Dec1 and Dec2 are regulators of the mammalian molecular clock. Nature 419, 841–844 (2002).

38. Cook, M. E., Jarjour, N. N., Lin, C.-C. & Edelson, B. T. Transcription Factor Bhlhe40 in Immunity and Autoimmunity. Trends Immunol. 41, 1023–1036 (2020).

39. Bennett, R. L. et al. A mutation in histone H2B represents a new class of oncogenic driver. Cancer Discov. 9, 1438–1451 (2019).

40. McGinty, R. K. & Tan, S. Nucleosome structure and function. Chem. Rev. 115, 2255–2273 (2015).

41. McBride, M. J. et al. The nucleosome acidic patch and H2A ubiquitination underlie mSWI/SNF recruitment in synovial sarcoma. Nat. Struct. Mol. Biol. 27, 836–845 (2020).

42. Dao, H. T., Dul, B. E., Dann, G. P., Liszczak, G. P. & Muir, T. W. A basic motif anchoring ISWI to nucleosome acidic patch regulates nucleosome spacing. Nat. Chem. Biol. 16, 134–142 (2020).

43. Skrajna, A. et al. Comprehensive nucleosome interactome screen establishes fundamental principles of nucleosome binding. Nucleic Acids Res. 48, 9415–9432 (2020).

44. Anink-Groenen, L. C. M., Maarleveld, T. R., Verschure, P. J. & Bruggeman, F. J. Mechanistic stochastic model of histone modification pattern formation. Epigenetics and Chromatin 7, 1–16 (2014).

45. Tachiwana, H. et al. Chromatin structure-dependent histone incorporation revealed by a genomewide deposition assay. Elife 10, 1–30 (2021).

46. Filippakopoulos, P. et al. Selective inhibition of BET bromodomains. Nature 468, 1067–1073 (2010).

47. Shi, J. & Vakoc, C. R. The Mechanisms behind the Therapeutic Activity of BET Bromodomain Inhibition. Mol. Cell 54, 728–736 (2014).

48. Savitski, M. M. et al. Multiplexed Proteome Dynamics Profiling Reveals Mechanisms Controlling Protein Homeostasis. Cell 173, 260–274.e25 (2018).

49. Stein, E. M. et al. The DOT1L inhibitor pinometostat reduces H3K79 methylation and has modest clinical activity in adult acute leukemia. Blood 131, 2662–2669 (2018).

50. Gilan, O. et al. Functional interdependence of BRD4 and DOT1L in MLL leukemia. Nat. Struct. Mol. Biol. 23, 673–681 (2016).

51. Osman, S. & Cramer, P. Structural Biology of RNA Polymerase II Transcription: 20 Years On. Annu. Rev. Cell Dev. Biol. 36, 1–34 (2020).

52. Cramer, P. Organization and regulation of gene transcription. Nature 573, 45–54 (2019).

53. Young, R. A. RNA POLYMERASE II. Annu. Rev. Biochem. 60, 689–715 (1991).

54. Santo, L. et al. AT7519, A novel small molecule multi-cyclin-dependent kinase inhibitor, induces apoptosis in multiple myeloma via GSK-3B activation and RNA polymerase II inhibition. Oncogene 29, 2325–2336 (2010).

55. Harlen, K. M. & Churchman, L. S. The code and beyond: Transcription regulation by the RNA polymerase II carboxy-terminal domain. Nat. Rev. Mol. Cell Biol. 18, 263–273 (2017).

