## Supplementary Information for "Tracking chromatin state changes using *μ*Map photo-proximity labeling"

### Supporting Information

**Ciaran P. Seath*, Antony J. Burton*, David W. C. MacMillan†, Tom W. Muir†**

[1]*Merck Center for Catalysis at Princeton University, Princeton, New Jersey 08544, USA.*

[2]*Department of Chemistry, Princeton University, Princeton, New Jersey 08544, USA*

*These authors contributed equally to this work.

<u>Supplementary Figures</u>

#### Supporting Figure 1 – Synthesis of Ir-Cfa$^{C}$

H-VKIISRKSLGTQNVYDIGVEKDHNFLLKNGLVASN**CF**GSG**K**G-$NH_2$

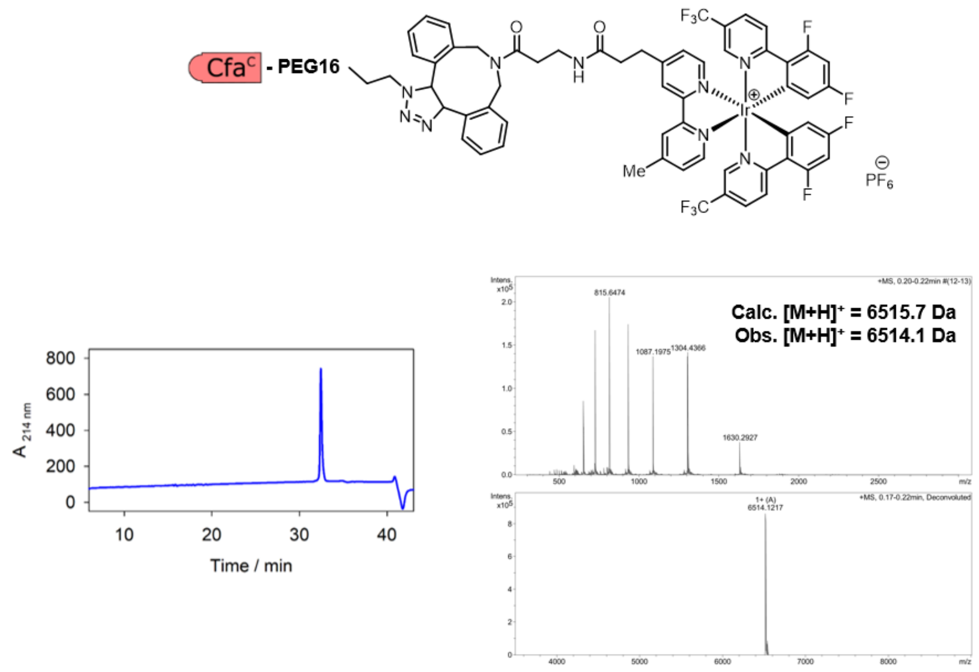

**Supporting Figure 1**. Sequence of Cfa$^{C}$ synthesized for use in this study. Extein residues are shown in bold, Ir photocatalyst is conjugated to the ε-amino group of the lysine highlighted in red. Also shown is the structure of the DBCO-conjugated Ir photocatalyst used in this study. RP-HPLC trace (left; 214 nm) and ESI-MS spectrum (right; raw and deconvoluted spectra) for purified Cfa$^{C}$-Ir.

#### Supporting Figure 2 –TMT-based proteomics for H3.1

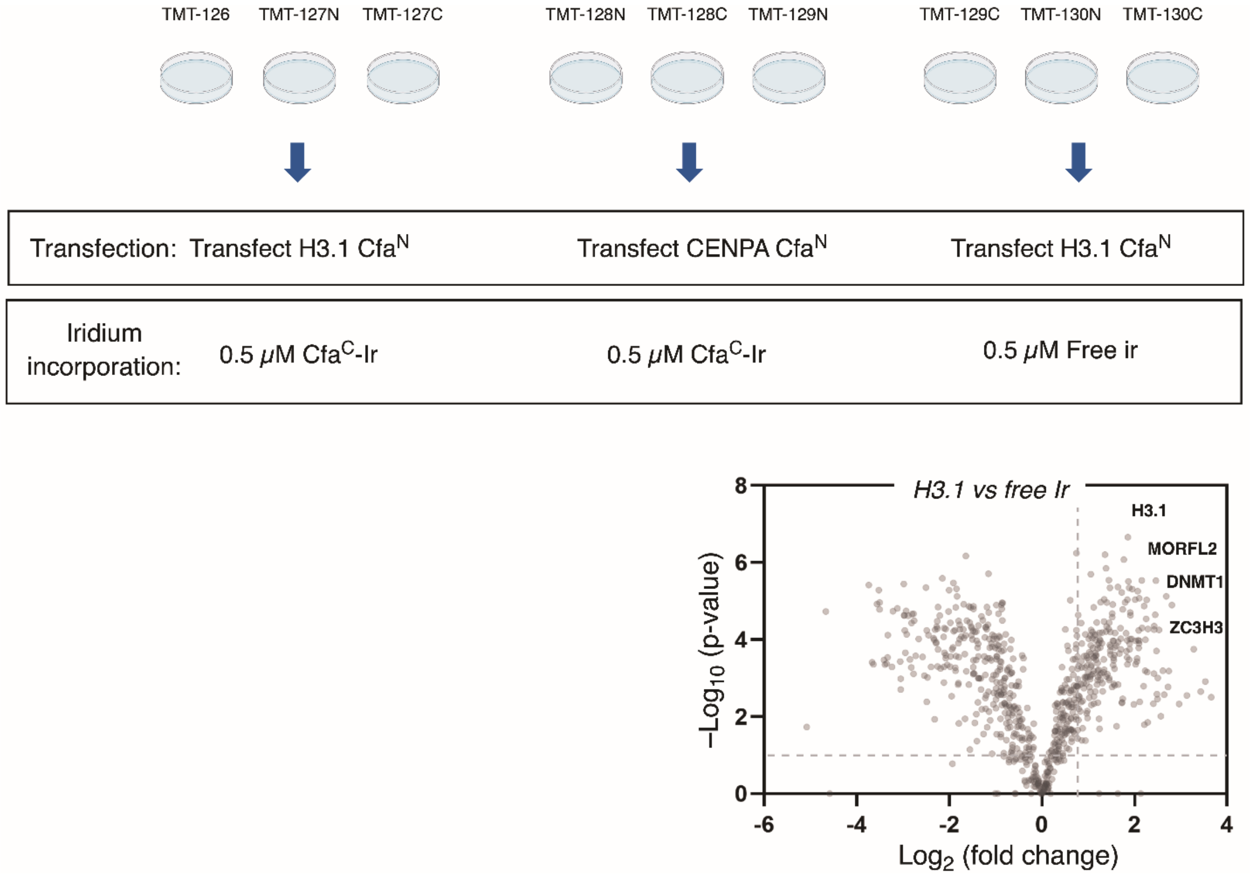

**Supporting Figure 2**. Top: TMT 10-plex setup for H3.1 and CENPA interactomics. Bottom: Volcano plot for H3.1 versus free Ir. Cut-offs = >0.5 $Log_2$-fold change, <0.05 FDR-corrected p value.

#### Supporting Figure 3 – Analysis of TMT-based proteomics for H3.1

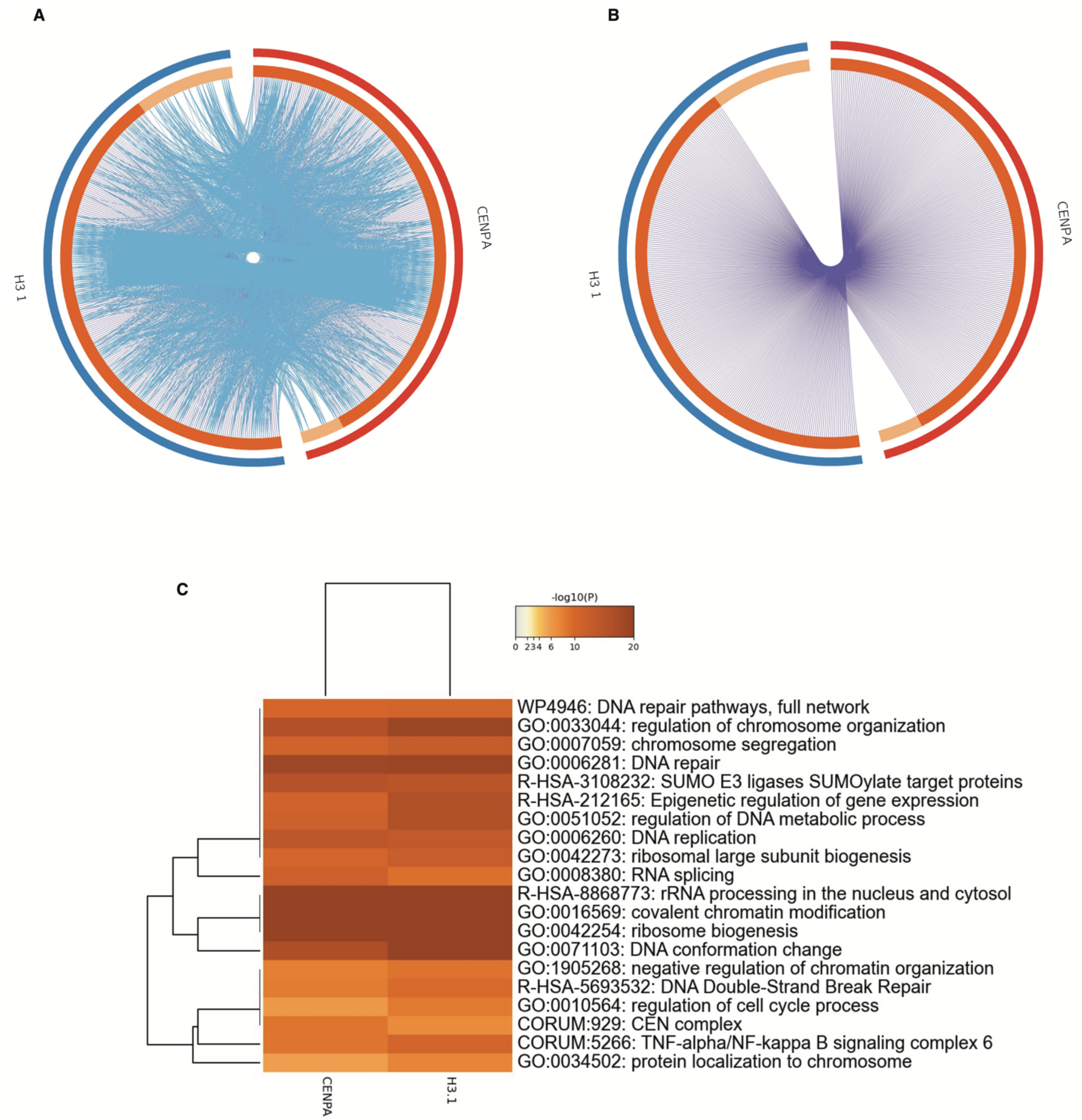

**Supporting Figure 3**. a) Circos plot showing overlap (pink lines) and functional relationships (blue lines) between H3.1 hits and CENPA hits. a) Circos plot showing overlap (pink lines) H3.1 hits and CENPA hits. c) Heatmap showing comparative enrichment of GO terms for H3.1 hits and CENPA hits. Cut-offs = >0.5 $Log_2$-fold change, <0.05 FDR-corrected p value.

#### Supporting Figure 4 – Incorporation and splicing blots for H2A and H2A E92K

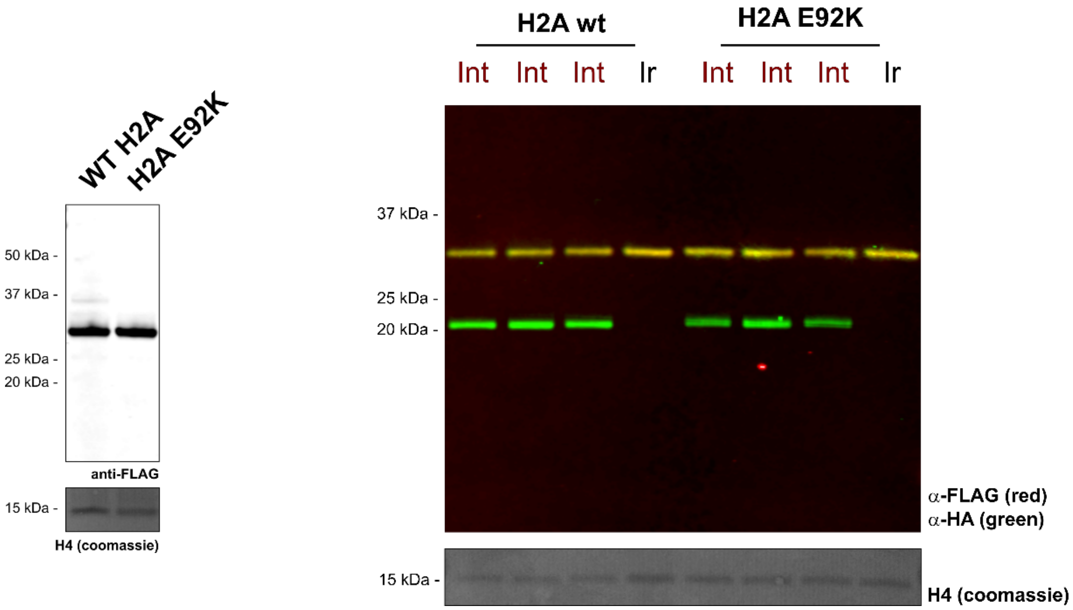

**Supporting Figure 4**. Left: anti-FLAG western blot for the expression of H2A-HA-$Cfa^{N}$-FLAG and H2A-E92K-HA-$Cfa^{N}$-FLAG. H4 visualized by coomassie blue stain is provided as a loading control. Right: In-nuclei splicing reactions for H2A and H2A E92K constructs in the presence of $Cfa^{C}$-Ir and free Ir as visualized by anti-HA and ant-FLAG western blot. H4 visualized by coomassie blue stain is provided as a loading control. MW of H2A-HA-Cfa-N-FLAG = 28,121 Da.

#### Supporting Figure 5 – Incorporation and splicing blots for H4

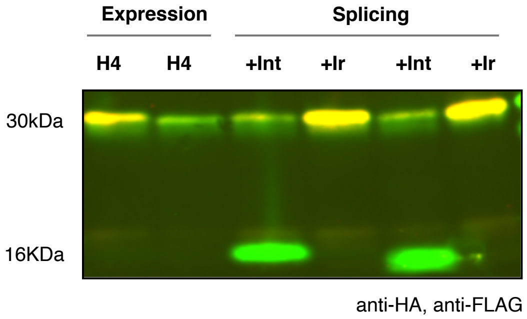

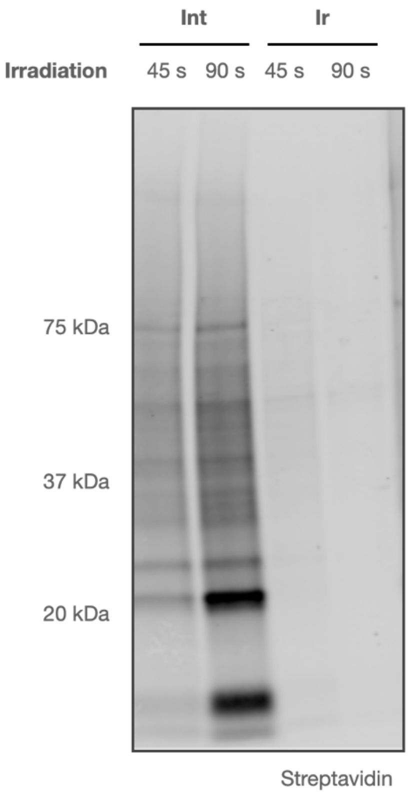

**Supporting Figure 5**. Left: anti-HA/anti-FLAG western blot visualizing the treatment of H4-HA-$Cfa^N$-FLAG with $Cfa^C$-Ir or free Ir. Right: Labeling of nuclear proteins after installation of Ir photocatalyst (45 and 90 second irradiation with blue LEDs) versus free Ir control as visualized by streptavidin-800. MW of H4-HA-Cfa-N-FLAG = 25,278 Da.

#### Supporting Figure 6– Blots for H1.1/1.3

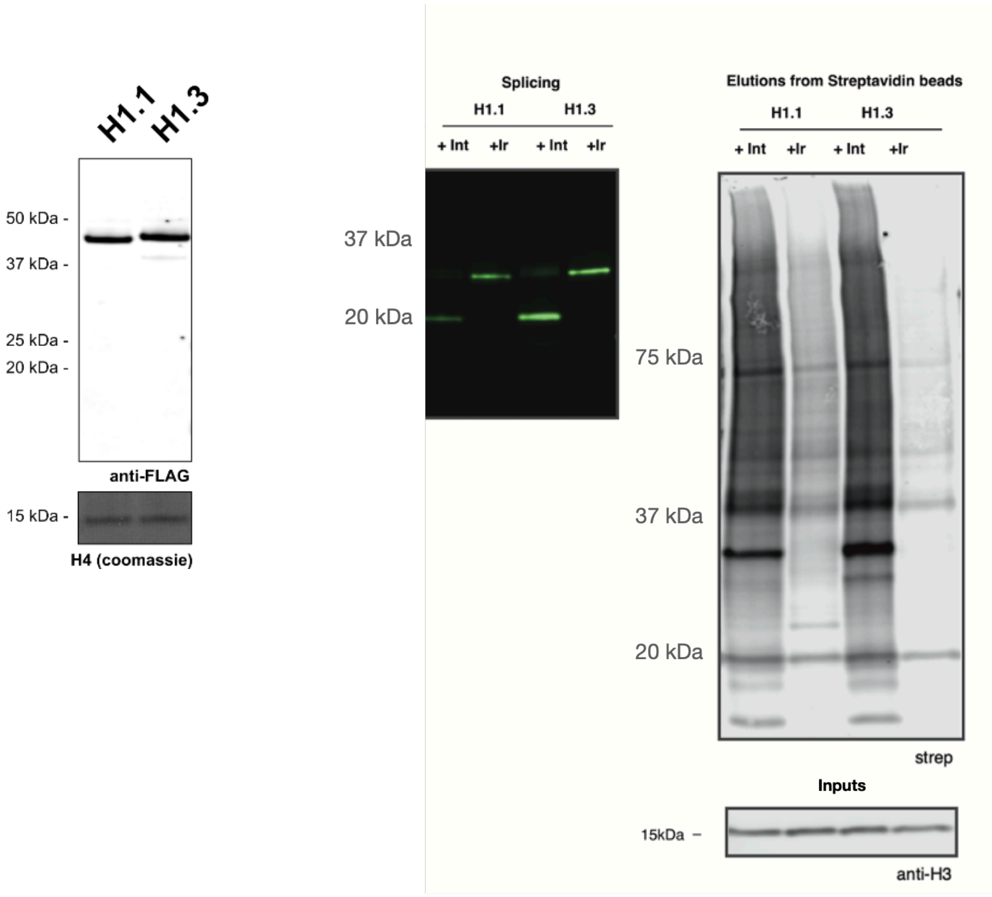

**Supporting Figure 6**. Left: anti-FLAG western blot for the expression of H1.1-HA-Cfa$^N$-FLAG and H1.3-HA-Cfa$^N$-FLAG. H4 visualized by coomassie blue stain is provided as a loading control. Middle: In-nuclei splicing reactions for H1.1 and H1.3 constructs in the presence of Cfa$^C$-Ir (left) and free Ir (right) as visualized by anti-HA western blot. Anti-H3 visualized by western blot is provided as a loading control. Right: Labeling of nuclear proteins after installation of Ir photocatalyst (left) versus free Ir control (right) as visualized by streptavidin-800. MW of H1.1-HA-Cfa-N-FLAG = 35,867 Da. MW of H1.3-HA-Cfa-N-FLAG = 36,375 Da.

#### Supporting Figure 7– TMT-based proteomics for H1

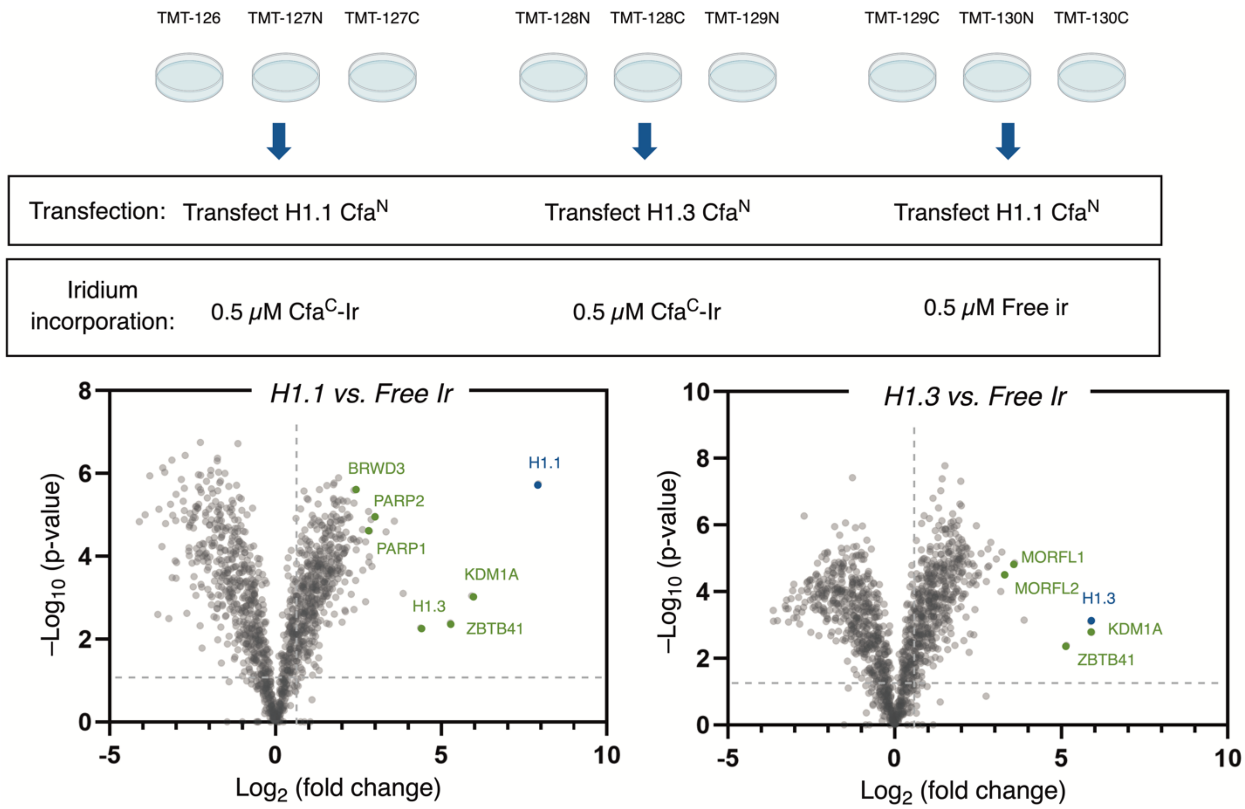

**Supporting Figure 7**. Top: TMT 10-plex setup for H1.1 and H1.3 interactomics. Bottom: Volcano plots for H1.1 (left) and H1.3 (right) versus free Ir. Cut-offs = >0.5 $Log_2$-fold change, <0.05 FDR-corrected p value.

#### Supporting Figure 8 – Comparative analysis of TMT-based proteomics H1.1 and H1.3

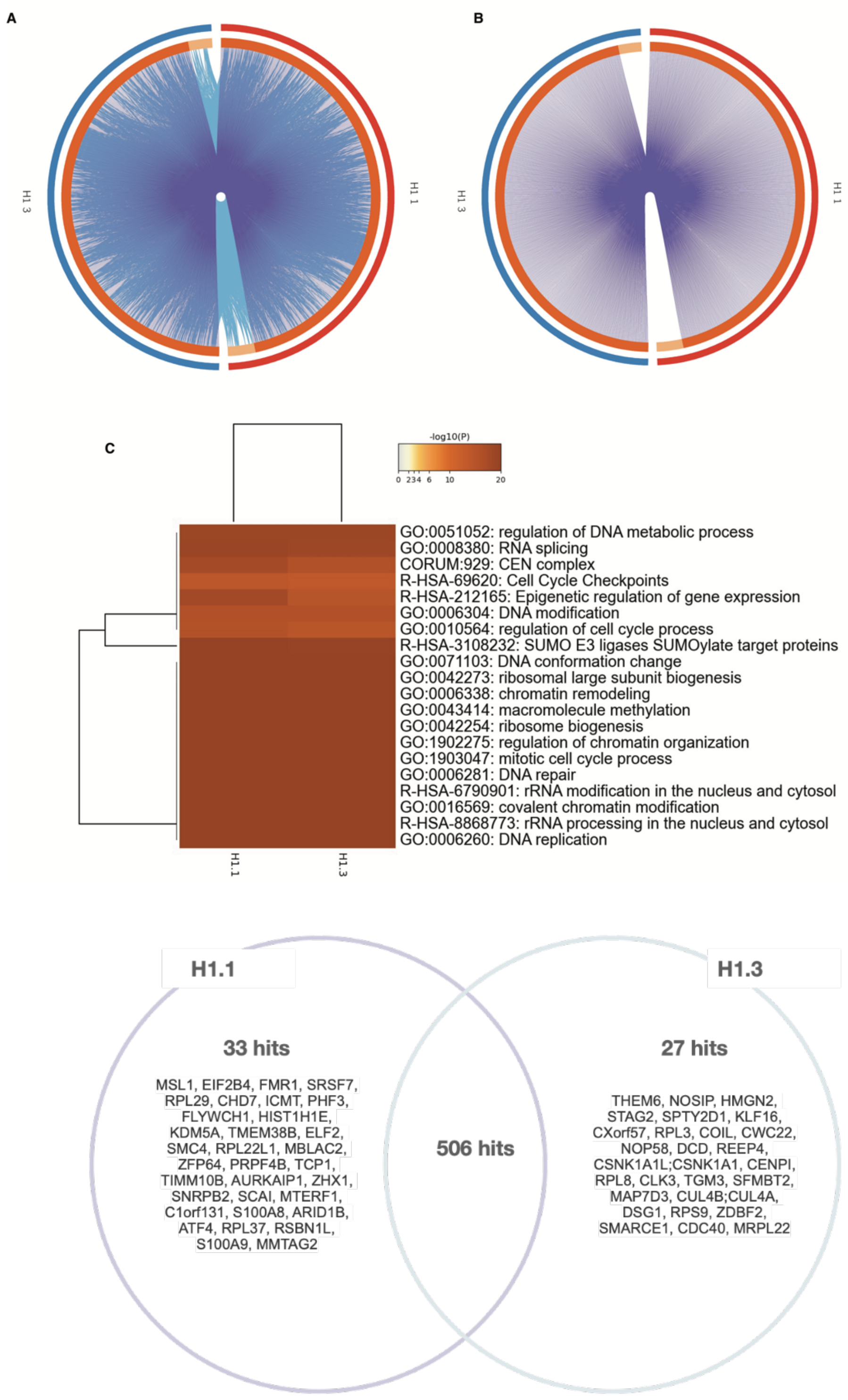

**Supporting Figure 8**. Top: a) Circos plot showing overlap (pink lines) and functional relationships (blue lines) between H1.1 hits and H1.3 hits. b) Circos plot showing overlap (pink lines) between H1.1 hits and H1.3 hits. c) Heatmap showing comparative enrichment of GO terms for H1.1 hits and H1.3 hits. Bottom: Venn diagram showing comparison of hits from H1.1 and H1.3 proteomics. Cut-offs = >$Log_2$-fold change, <0.05 FDR-corrected p value.

#### Supporting Figure 9 – Comparison of H1.1 hits to previous datasets

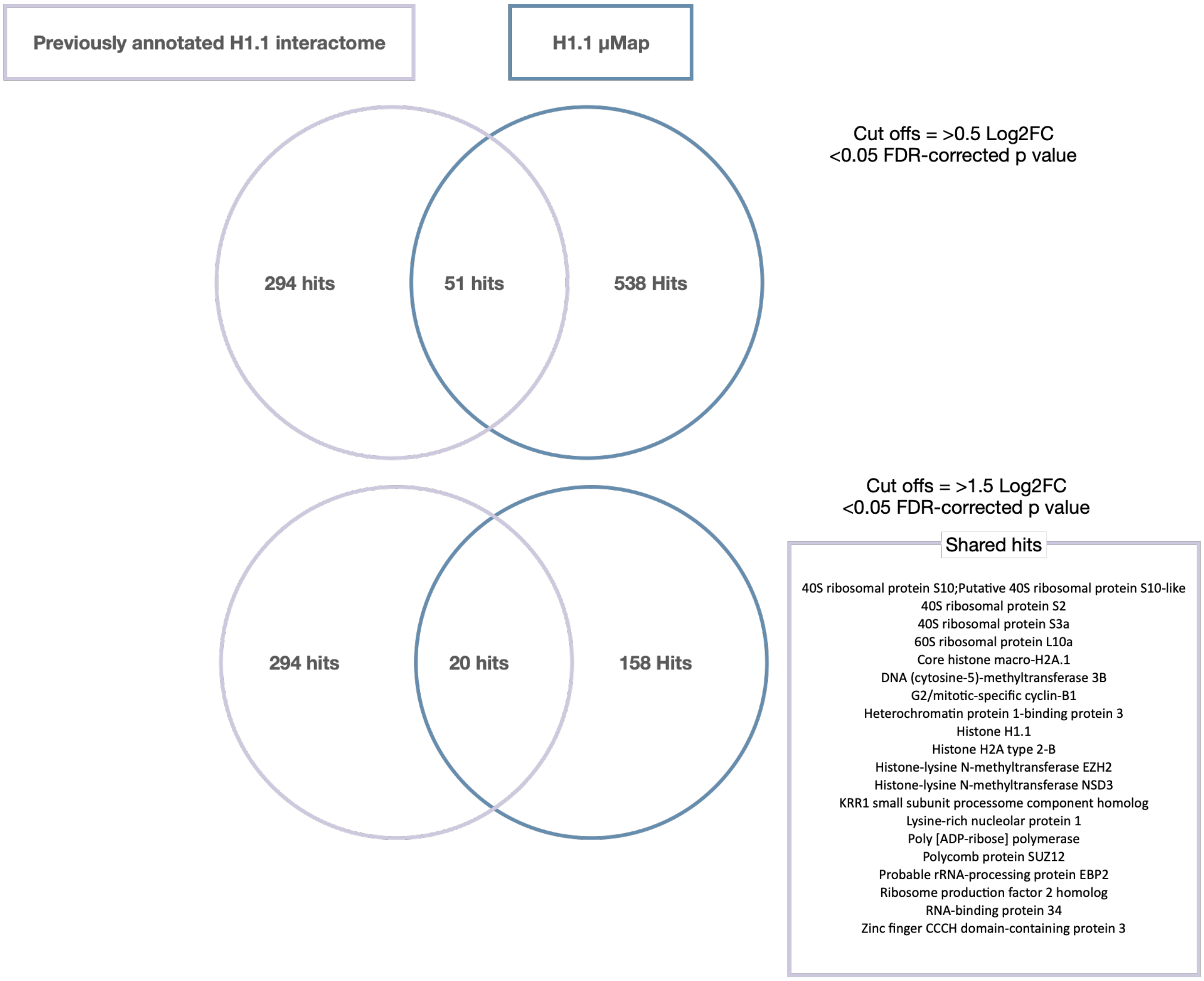

**Supporting Figure 9**. Top: Comparison of hits from ComPPI database (https://comppi.linkgroup.hu/protein_search/interactors/Q02539) with hits from H1.1 µMap using >0.5 Log2FC cut off. This shows 51 overlapping proteins across datasets. Bottom: Comparison of hits from ComPPI database (https://comppi.linkgroup.hu/protein_search/interactors/Q02539) with hits from H1.1 µMap using >1.5 Log2FC cut off. This shows 20 overlapping proteins across datasets. Shared hits are listed.

#### Supporting Figure 10 – Distribution analysis of H1.1/H2A/H3.1/H4 proteomics

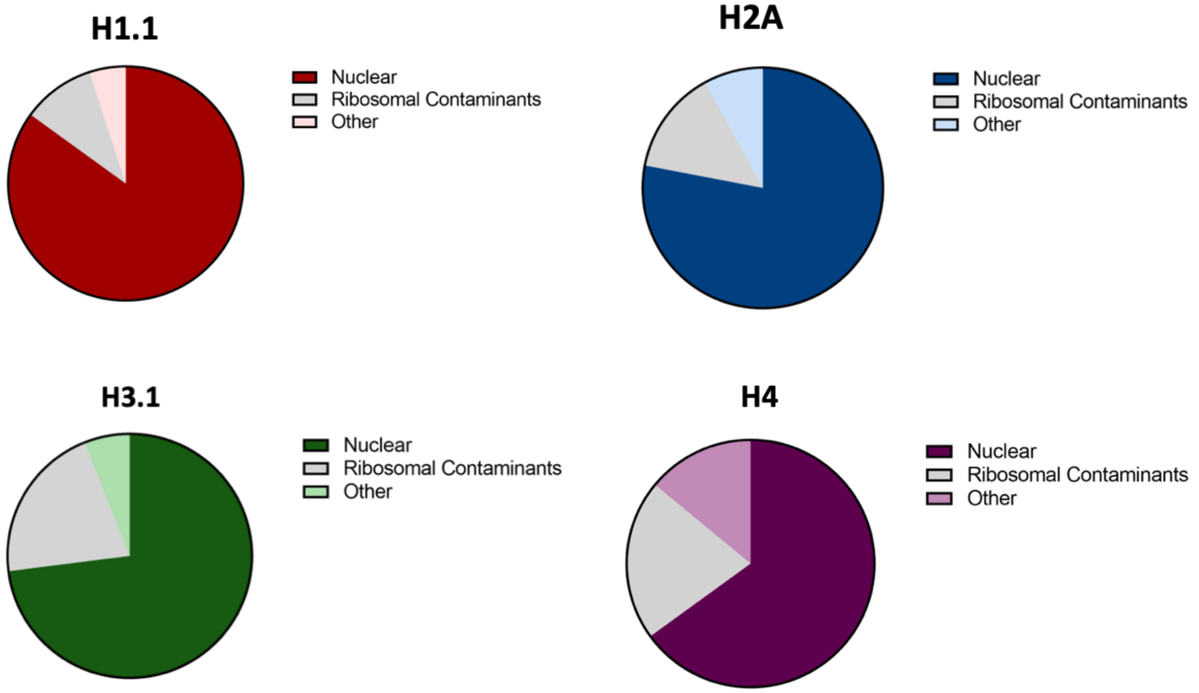

**Supporting Figure 10**. Top left: Pie chart showing the distribution of hits in H1.1 (N = 158). Top right: Pie chart showing the distribution of hits in H2A (N=120). Bottom left: Pie chart showing the distribution of hits in H3.1 (N=100). Bottom right: Pie chart showing the distribution of hits in H4 (N=81). Cut-offs = >1.5 $Log_2$-fold change, <0.05 FDR-corrected p value. Ribosomal contaminants made up of 40S and 60S ribosomal proteins.

#### Supporting Figure 11 – Iridium is localized at the DNA entry/exit site on H2A

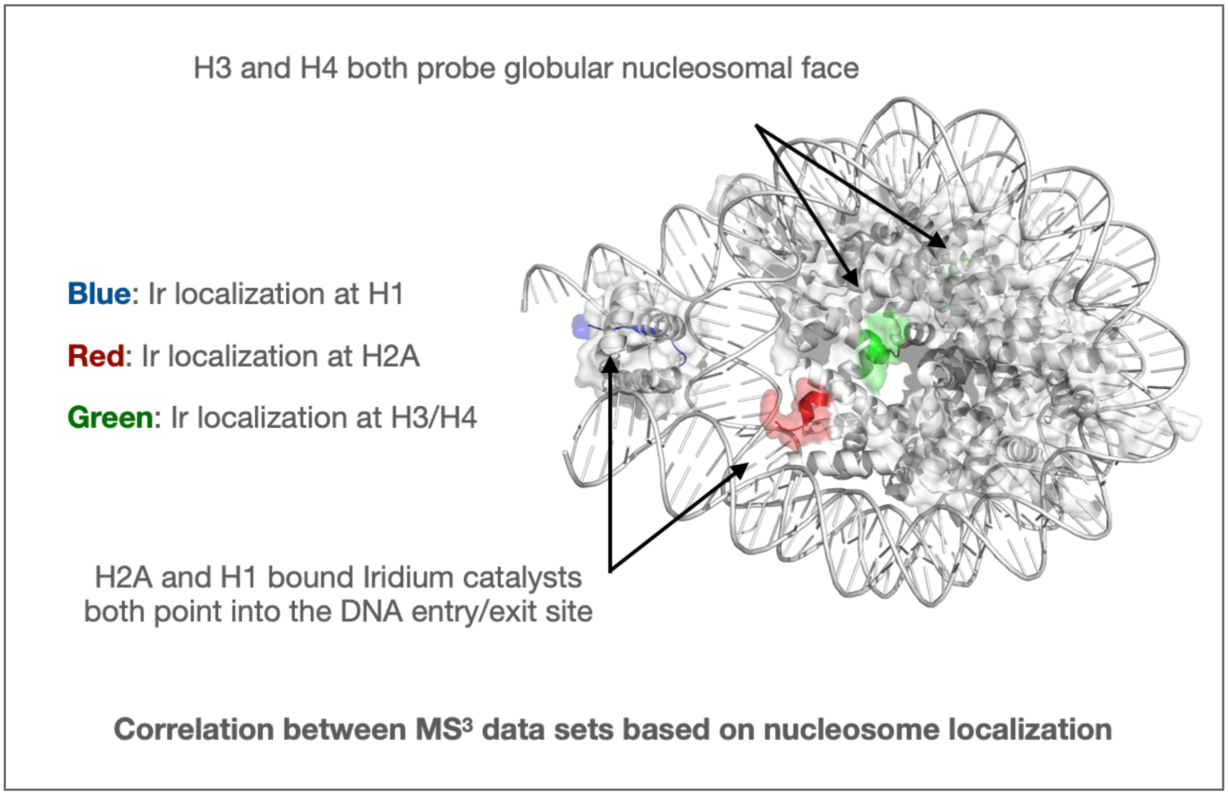

**Supporting Figure 11**. PyMol image of the chromatosome (7k5x) showing approximate nucleosomal locations of Ir-catalyst incorporation. Correlation analysis shows significant correlations between H2A and H1 hits and H3/H4 hits (shown in heatmap in Figure 2g). Analysis performed using Heatmapper.ca. Euclidean hierarchical clustering calculated as average linkage.

#### Supporting Figure 12 – CTCF/SMC1A expression and splicing

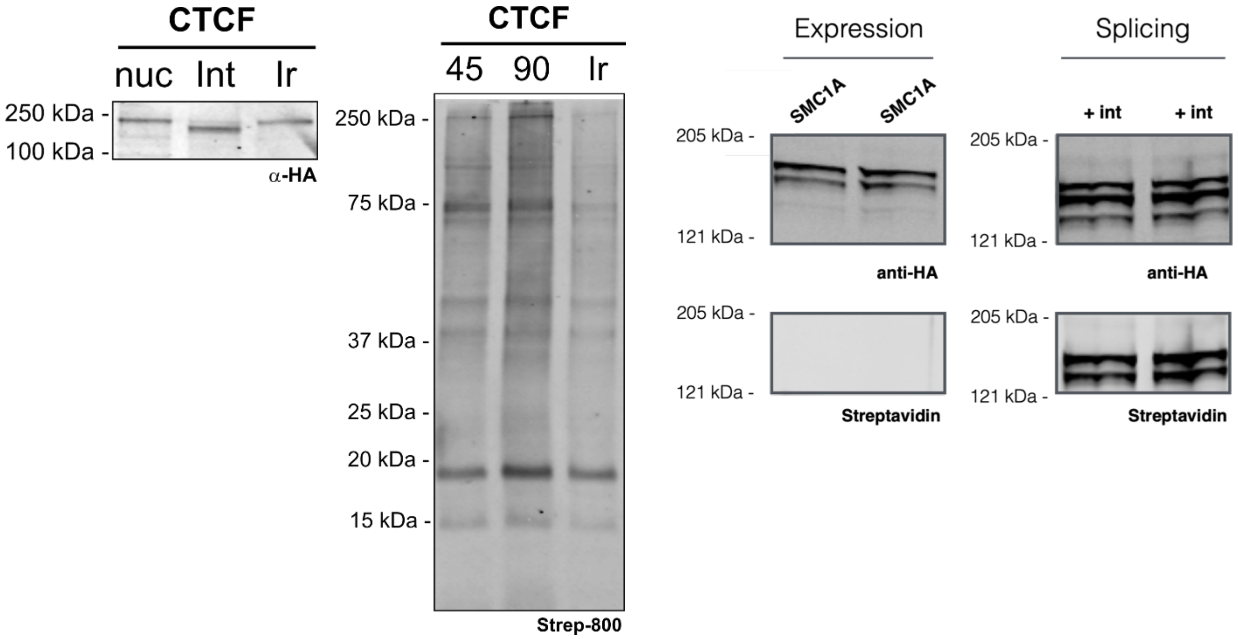

**Supporting Figure 13**. Left: anti-HA western blot visualizing the treatment of CTCF-HA-Cfa$^{N}$-FLAG with Cfa$^{C}$-Ir and free Ir catalyst (nuc = lysed nuclei; Int = treatment with Cfa$^{C}$-Ir; Ir = treatment with free Ir catalyst). Middle: Labeling of nuclear proteins after installation of Ir photocatalyst (45 and 90 second irradiation with blue LEDs) versus free Ir control as visualized by streptavidin-800. Right: anti-HA and streptavidin western blots visualizing the SMC1A-HA-Cfa$^{N}$-FLAG with Cfa$^{C}$-biotin. MW of CTCF-HA-Cfa-N-FLAG = 96,811 Da. MW of SMC1A - HA-Cfa-N-FLAG = 157,258 Da.

#### Supporting Figure 13 – CTCF/SMC1A proteomics experiment

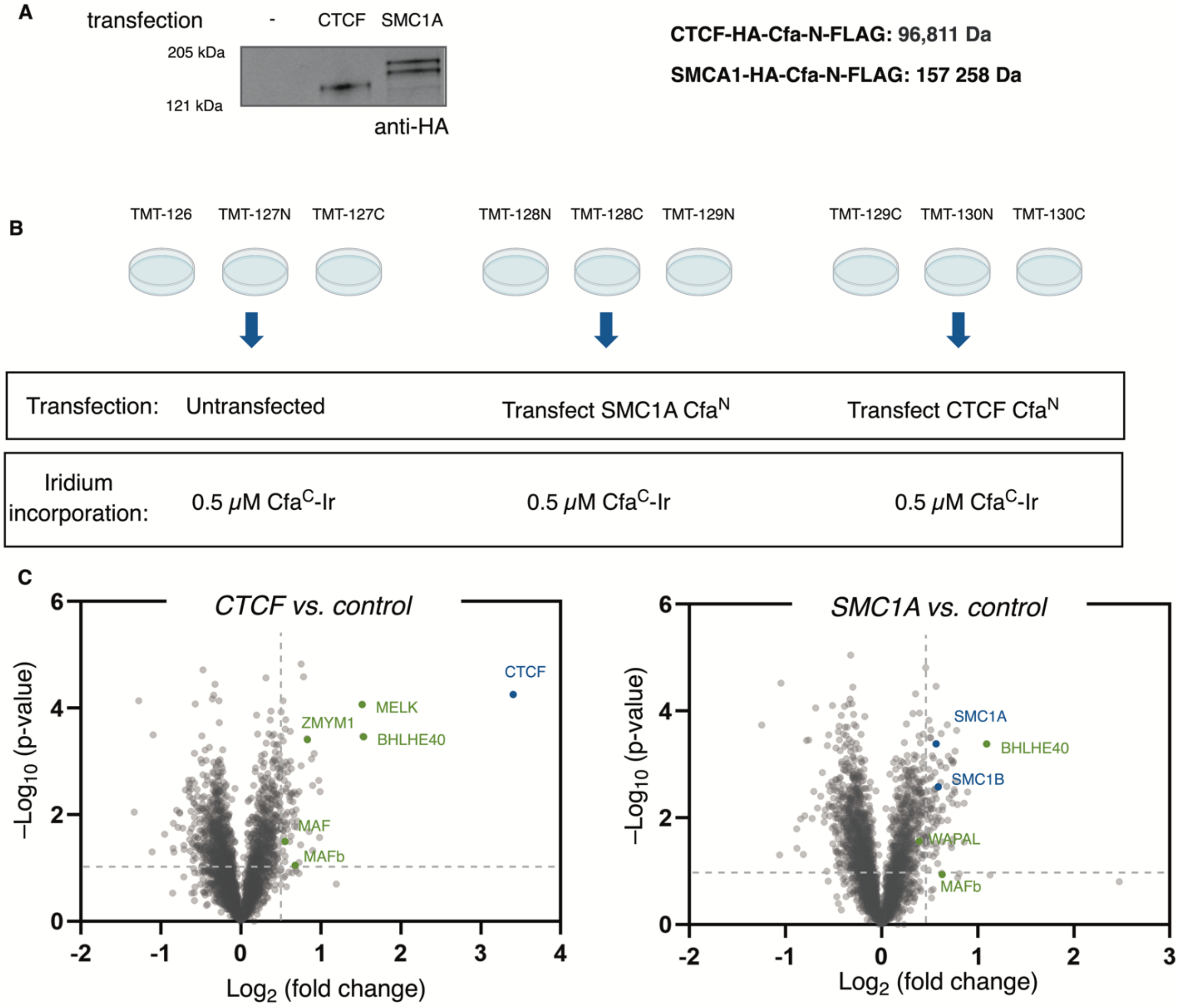

**Supporting Figure 13**. a) anti-HA western blot showing expression of CTCF-HA-Cfa$^{N}$-FLAG and SMC1A-HA-Cfa$^{N}$-FLAG. b) TMT 10-plex setup for SMC1A and CTCF interactomics. c) Volcano plots for CTCF vs free Ir (left) and SMC1A vs free Ir (right). Cut-offs = >0.5 Log$_2$-fold change, <0.05 FDR-corrected p value.

#### Supporting Figure 14 – CTCF/SMC1A GO analysis

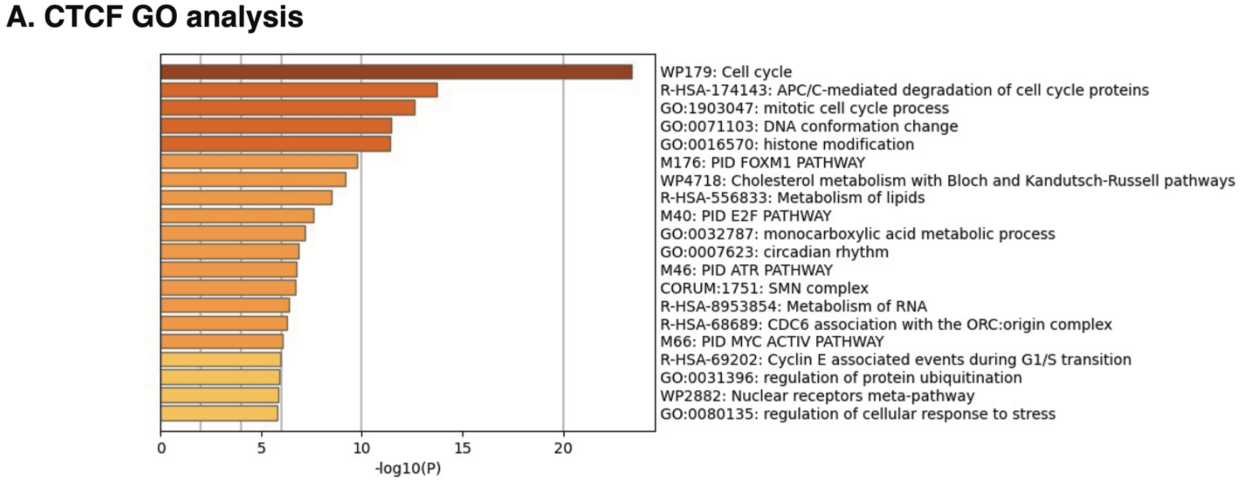

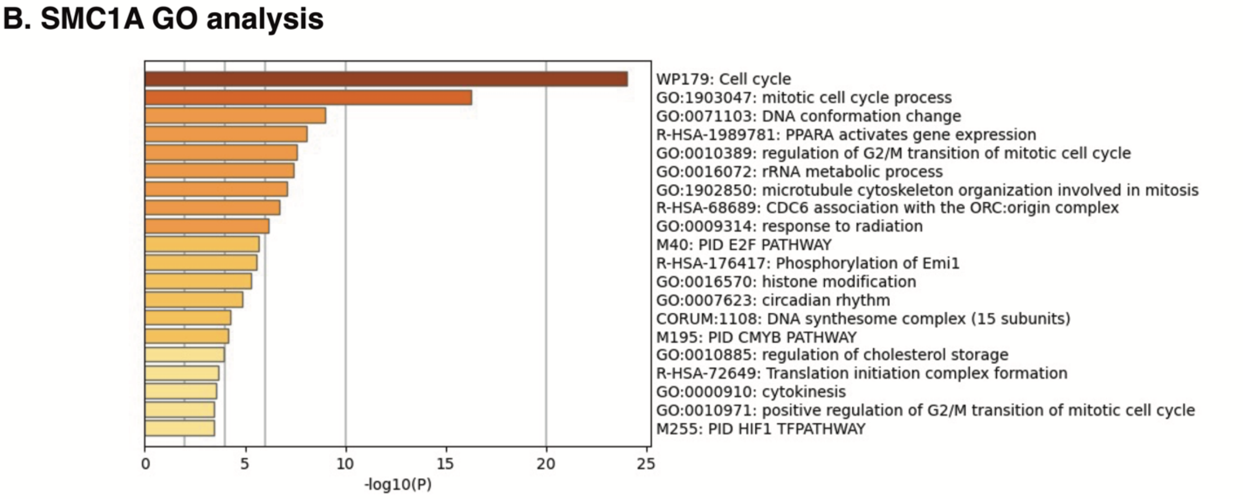

**Supporting Figure 14**. a) Top GO terms enriched in analysis of CTCF hits. b) Top GO terms enriched in analysis of SMC1A hits. Cut-offs = >0.5 $Log_2$-fold change, <0.05 FDR-corrected p value.

### Supporting Figure 15 – SMC1A/CTCF combined GO and enrichment analysis

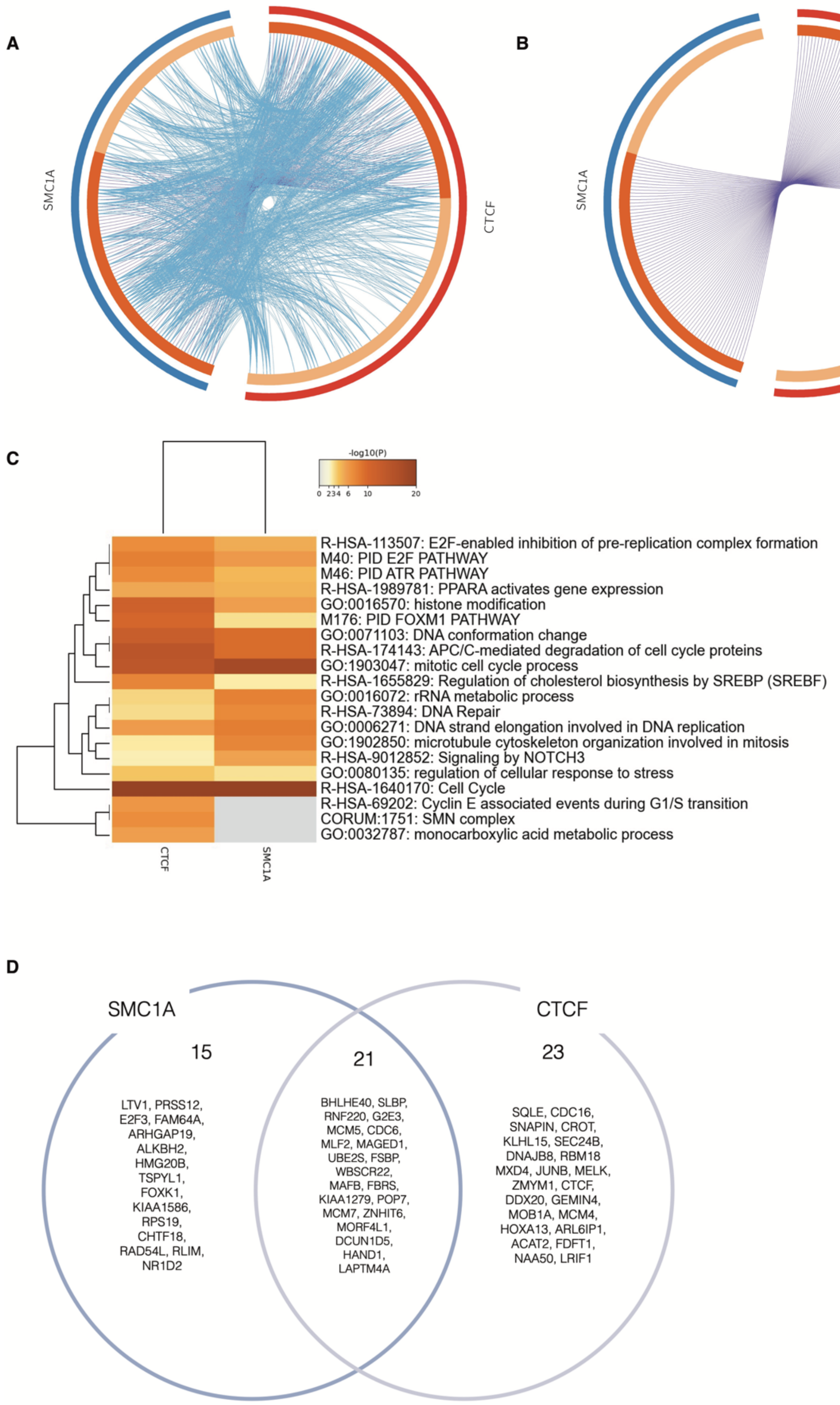

**Supporting Figure 15**. a) Circos plot showing overlap (pink lines) and functional relationships (blue lines) between CTCF hits and SMC1A hits. b) Circos plot showing overlap (pink lines) between CTCF hits and SMC1A hits. c) Heatmap showing comparative enrichment of GO terms for CTCF hits and SMC1A hits. d) Venn diagram showing overlapping and unique hits for CTCF hits and SMC1A hits. Cut-offs = >0.5 $Log_2$-fold change, <0.05 FDR-corrected p value.

#### Supporting Figure 16 – H2A with JQ1 and Pinometostat treatment splicing blot

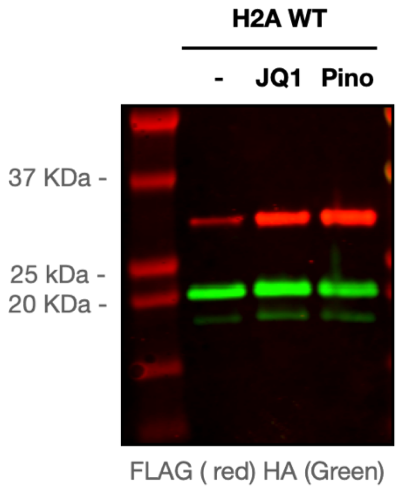

**Supporting Figure 16.** *In nucleo* splicing reactions for H2A constructs in the presence of Cfa$^{C}$-Ir as visualized by anti-HA/anti-FLAG western blot. Left (untreated cells), middle (cells treated with JQ1), right (cells treated with Pinometostat). MW of H2A-HA-Cfa-N-FLAG = 28,121 Da.

#### Supporting Figure 17 – H3K79(me)2 depletion in the presence of Pinometostat

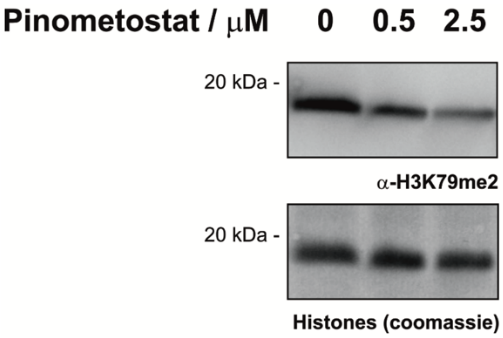

**Supporting Figure 17**. Treatment of HEK 293T cells with pinometostat for 24 h (0, 0.5, 2.5 μM) results in ≈60% decrease in H3K79 dimethylation, as visualized by western blotting with an anti-H3K79me2 antibody. Histones H3, H2A, and H2B, stained by coomassie blue, are provided as a loading control.

#### Supporting Figure 18 – CDK inhibitor treatment depletes Pol II CTD S2/5 phos

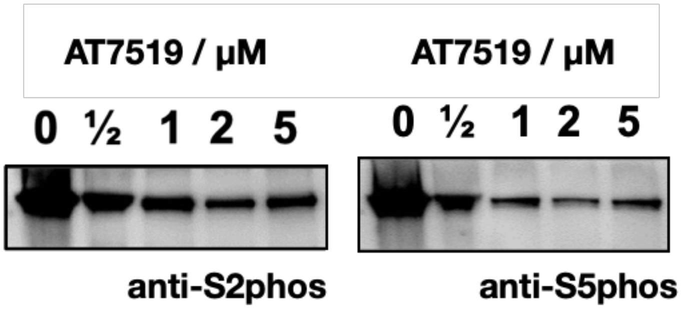

**Supporting Figure 18**. Treatment of HEK 293T cells with AT7519 for 2 h results in a decrease in CTD S2 and S5 phosphorylation, as visualized by western blotting with anti-S2phos and anti-S5phos antibodies.

#### Supporting Figure 19 – Validation of RBP1 expression and splicing

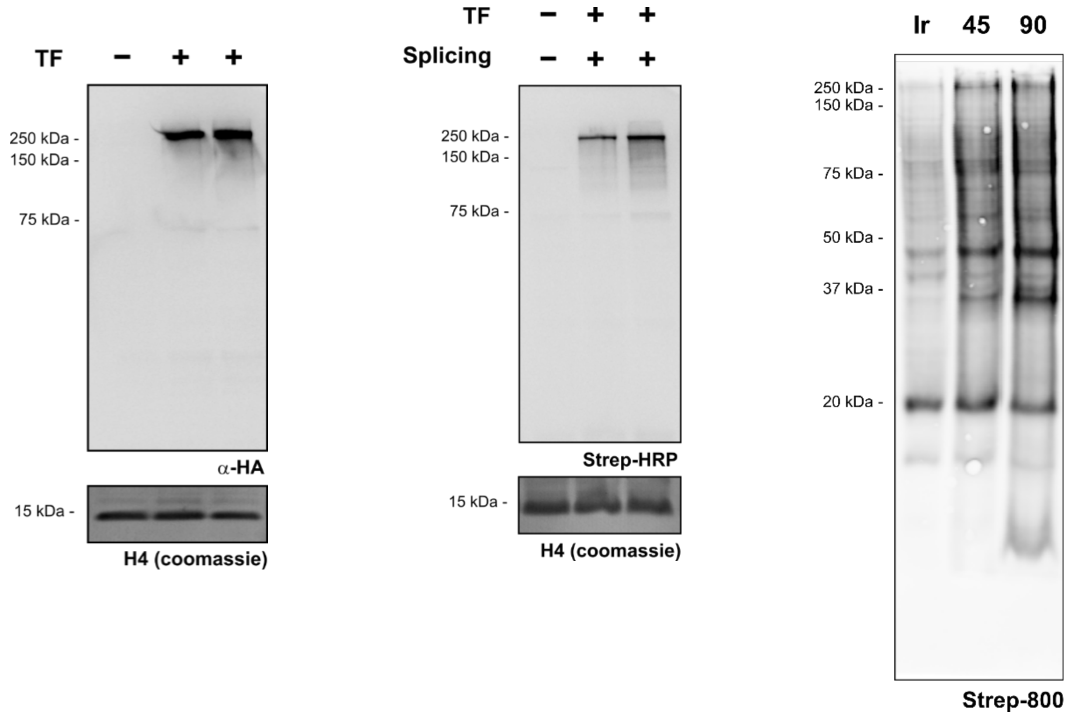

**Supporting Figure 19**. Left: Anti-HA western blot for the expression of RBP1-HA-Cfa$^{N}$-FLAG. H4 visualized by coomassie blue stain is provided as a loading control. Middle: *In nucleo* splicing reactions for RBP1-HA-Cfa$^{N}$-FLAG in the presence of free Ir (-) Cfa$^{C}$-biotin (+) as visualized by streptavidin-800. H4 visualized by coomassie blue stain is provided as a loading control. Right: Labeling of nuclear proteins after installation of Ir photocatalyst (45 and 90 second irradiation with blue LEDs) versus free Ir control as visualized by streptavidin-800. MW of RBP1-HA-Cfa$^{N}$-FLAG = 231,290 Da.

#### Supporting Figure 20 – RPB1 proteomics experiment

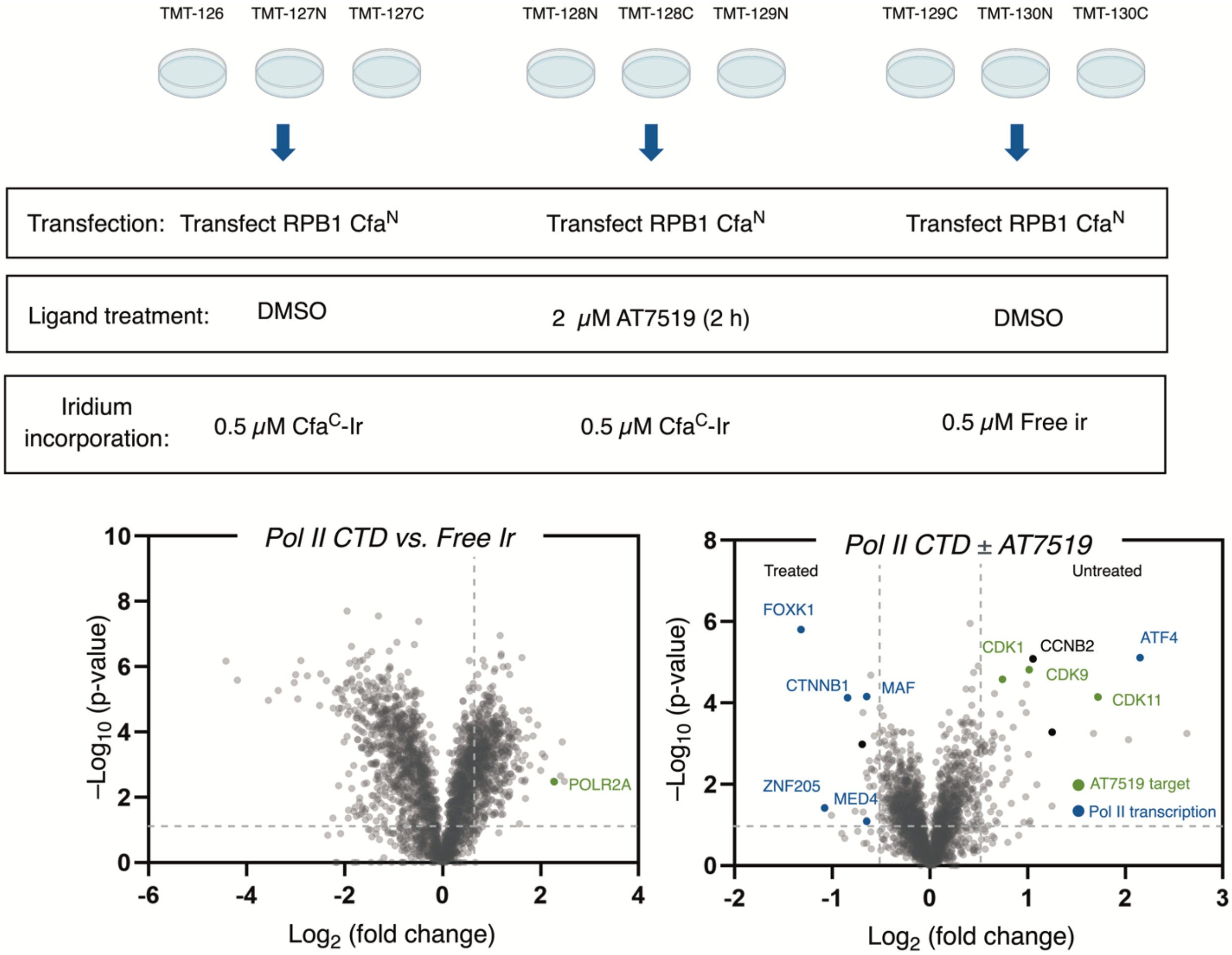

**Supporting Figure 20**. TMT 10-plex setup for RBP1 interactomics, in the presence and absence of 2 μM AT7519 (2 h pre-treatment). Volcano plots for RPB1 vs free Ir (left) and RBP1 +/- AT7519 (right). Cut-offs = >0.5 $Log_2$-fold change, <0.05 FDR-corrected p value.

#### Supporting Figure 21 – GO analysis Pol II

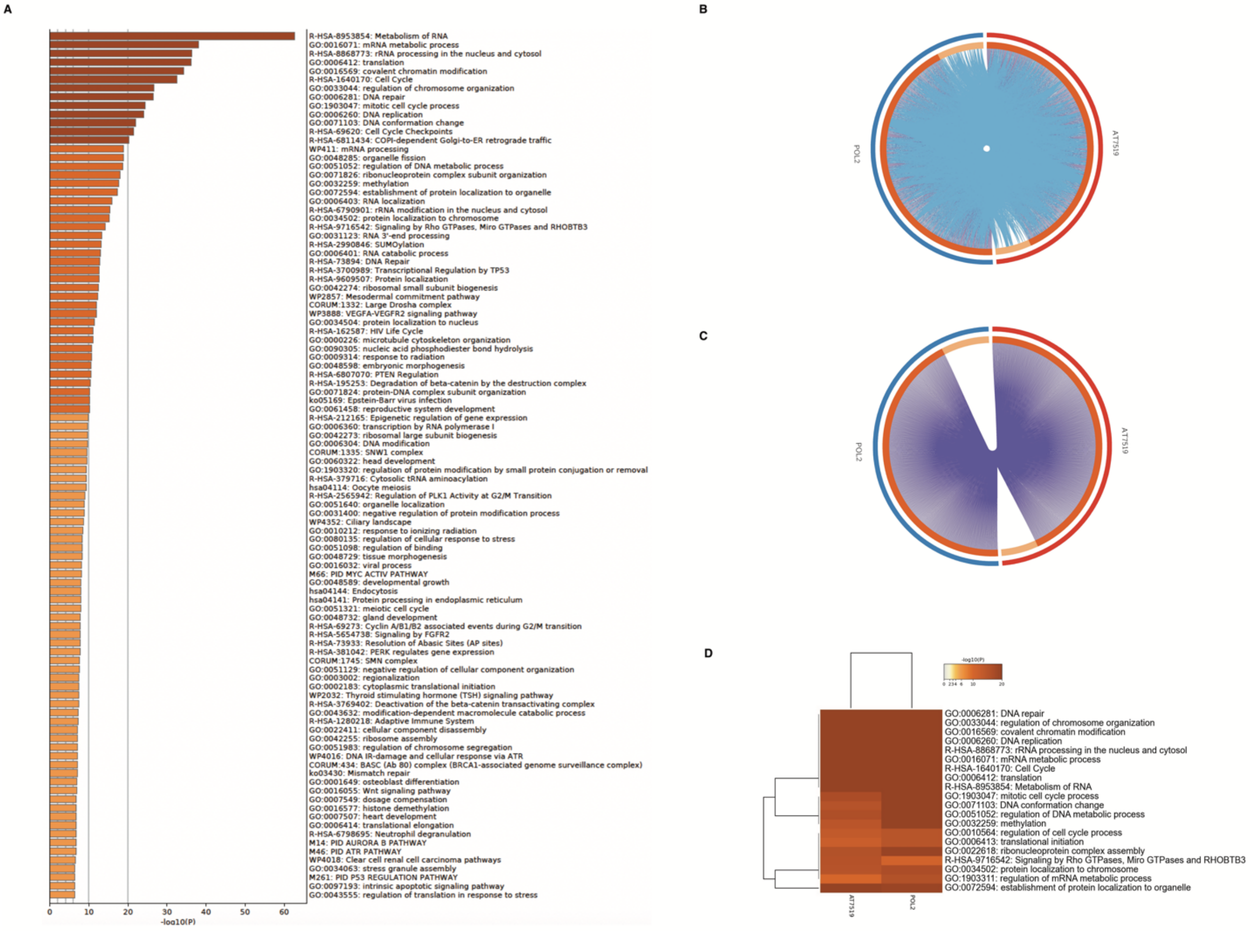

**Supporting Figure 21**. a) Full GO analysis of RPB1 hits. b) Circos plot showing overlap (pink lines) and functional relationships (blue lines) between RPB1 hits with and without AT7519 treatment. c) Circos plot showing overlap (pink lines) of RPB1 hits with and without AT7519 treatment. d) Heatmap showing comparative enrichment of GO terms for RPB1 hits with and without AT7519 treatment. Cut-offs = >0.5 $Log_2$-fold change, <0.05 FDR-corrected p value.

#### Supporting Figure 22 – Synthesis of Cfa$^{C}$- biotin

H-VKIISRKSLGTQNVYDIGVEKDHNFLLKNGLVASN**CF**GSG**K**G-$NH_2$

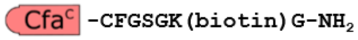

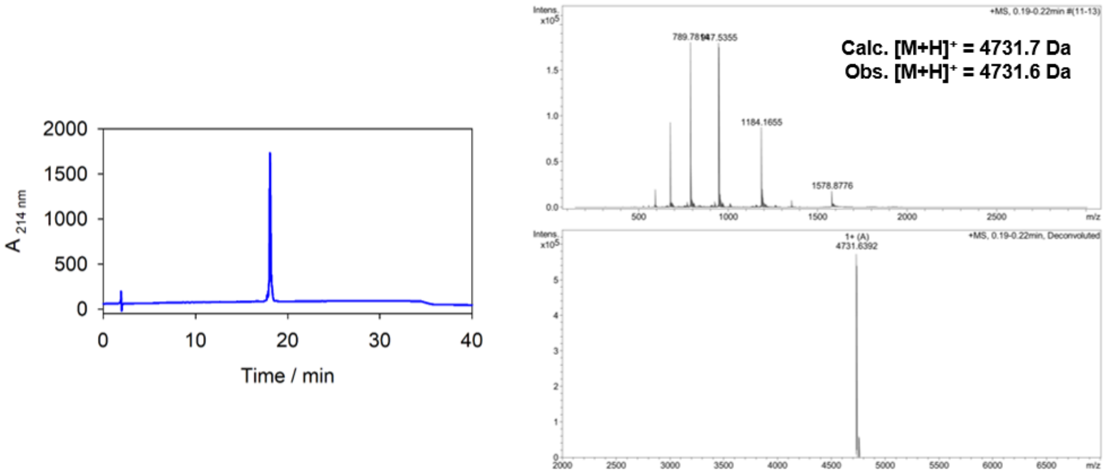

**Supporting Figure 22**. Sequence of Cfa$^{C}$-biotin synthesized for use in this study. Extein residues are shown in bold, biotin is conjugated to the ε-amino group of the lysine highlighted in red. RP-HPLC trace (left; 214 nm) and ESI-MS spectrum (right; raw and deconvoluted spectra) for purified Cfa$^{C}$-biotin.

#### General Considerations

Organic solvents were purified according to the method of Grubbs[1]. Water was purified using a Millipore Milli-Q Integral Water Purification System. Organic solutions were concentrated under reduced pressure on a Büchi rotary evaporator using a water bath. $^{1}$H NMR spectra were recorded on a Bruker UltraShield Plus Avance III 500 MHz unless otherwise noted and are internally referenced to residual solvent signals. Data for $^{1}$H NMR are reported as follows: chemical shift (δ ppm), multiplicity (s = singlet, d = doublet, t = triplet, q = quartet, p = quintet, m = multiplet, dd = doublet of doublets, dt = doublet of triplets…etc, br = broad), coupling constant (Hz) and integration. $^{13}$C NMR spectra were recorded on a Bruker UltraShield Plus Avance III 500 MHz (125 MHz) and data are reported relative to the solvent employed. High resolution mass spectra and intact protein mass spectra were obtained from the Princeton University Mass Spectral Facility and Princeton Proteomics & Mass Spectrometry Core. Irradiation of samples was performed in a PennOC Photoreactor. Chromatographic purification was carried out using a Biotage Isolera Prime flash chromatography system with SilaSep flash cartridges (60 mesh) and UV detection. Fmoc-protected amino acids were purchased form Matrix Innovations (Quebec, Canada). Diisopropylcarbodiimide (DIC) and Oxyma were purchased from Advanced Chemtech (Louisville, KY) and Chem-Impex (Wood Dale, IL), respectively. H-Rink Amide resin was obtained from Biotage (Charlotte, NC). Trifluoroacetic acid (TFA) was purchased from Halocarbon (North Augusta, SC). (7-Azabenzotriazol-1-yloxy)tripyrrolidinophosphonium hexafluorophosphate (PyAOP) was purchased from Matrix Innovations (Quebec, Canada). All buffers and synthetic starting materials were used as received from commercial sources. Bovine serum albumin (BSA) (A7906), Eppendorf Protein LoBind tubes (Z666505), Coppe(II) sulfate pentahydrate (7758-99-8) were purchased from Millipore Sigma (St. Louis, MO). 1X DPBS (14190144), Pierce BCA Protein Assay Kit (23227), and iBright Prestained Protein ladder (LC5615) were purchased from Thermo Scientific (Rockford, IL). TBST (IBB-581X) was purchased from Boston BioProducts (Ashland, MA). 5M Sodium chloride (S24600-500.0) was purchased from Research Products International (Mt. Prospect, S19 IL). Criterion TGX precast gels (5671044) and 4x Laemmli sample buffer (161-0747) were purchased from Bio-Rad (Hercules, CA). 20% SDS solution (351-066-721) was purchased from Quality Biological (Gaithersburg, MD). Biotin-PEG3-diazirine was synthesized as described previously.

#### Antibodies used in this study.

| | | |
|---|---|---|
| Anti-FLAG | Sigma Aldrich (M2) | 1:1,000 in TBS-T |
| Anti-HA | Abcam (ab9110) | 1:1,000 in TBS-T |
| Anti-pan-Acetyl | Abcam (ab21623) | 1:1,000 in TBS-T |
| Anti-H3 | Abcam (ab1791) | 1:5,000 in TBS-T |
| Anti-H4 | Abcam (ab31830) | 1:1,000 in TBS-T |
| Anti-H3K79me2 | Abcam (ab3594) | 1:1,000 in TBS-T |
| Anti-RBP1-S2phos | Abcam (ab5095) | 1:500 in TBS-T |
| Anti-RBP1-S5phos | Abcam (ab5408) | 1:500 in TBS-T |
| Anti-Mouse | LI-COR IRDye 680 nm | 1:10,000 in TBS-T |
| Anti-Rabbit | LI-COR IRDye 800 nm | 1:10,000 in TBS-T |
| Streptavidin | LI-COR IRDye Strep-800 nm | 1:10,000 in TBS-T |
| Anti-Rabbit HRP conjugate | Bio-Rad (1706515) | 1:3,000 in TBS-T |

#### Solid Phase Peptide Synthesis

Boc-$N^{\alpha}$-$Cfa^{C}$-CFGSGK(alloc)G-$NH_2$ was synthesized on a 0.1 mmol scale by standard Fmoc solid-phase peptide synthesis using DIC-Oxyma activation on a CEM Liberty Blue microwave-assisted peptide synthesizer on ChemMatrix Rink amide resin. Each residue was double coupled during the synthesis. Fmoc deprotection was performed at room temperature by the addition of 20% piperidine in DMF with 0.1 M HOBt.

Alloc deprotection was performed by the addition of 0.1 eq. $Pd(PPh_3)_4$ and 2.5 eq. N,N'-dimethylbarbituric acid in DCM with nitrogen agitation. Treatment was performed twice at rt for 30 min. The resin was sequentially washed with 3x DCM, 3x DCM:DMF (1:1 v/v), 3x DMF, and 1x 5% w/v sodium diethyldithiocarbamate in DMF.

The resin was then split into 0.02 mmol aliquots and treated as follows:

1. $Cfa^{C}$-biotin

Biotin (5 eq.) was coupled to the deprotected lysine side chain with PyAOP (4.95 eq.) and DIEA (10 eq.) activation in NMP for 4 h with nitrogen agitation. The resin was washed with 3x NMP, 3x DMF, and 3x DCM. Side-chain deprotection and cleavage from the resin was affected by addition of a 92.5:2.5:2.5:2.5 v/v/v solution of TFA:TIPS:EDT:$H_2O$ for 130 minutes at rt. The cleavage solution was reduced to <5 mL volume under a positive pressure of $N_2$, and the crude peptide was precipitated using cold diethyl ether. The crude peptide was isolated by refrigerated centrifugation, resuspended in 50/50 v/v $H_2O$:MeCN with 0.1% TFA, and lyophilized to yield a white solid.

2. $Cfa^C$-Ir

The iridium photocatalyst was conjugated to the deprotected lysine side chain by treatment with NHS-Ir (1.2 eq.) and DIEA (2 eq.) in DMF for 2 h with nitrogen agitation in the absence of light. The resin was washed with 3x DMF, and 3x DCM. Side-chain deprotection and cleavage from the resin was affected by addition of a 95:2.5:2.5 v/v/v solution of TFA:TIPS:$H_2O$ for 130 minutes at rt in the absence of light. The cleavage solution was reduced to <5 mL volume under a positive pressure of $N_2$, and the crude peptide was precipitated using cold diethyl ether. The crude peptide was isolated by refrigerated centrifugation, resuspended in 50/50 v/v $H_2O$:MeCN with 0.1% TFA, and lyophilized to yield a pale yellow solid.

### HPLC purification

Semi-preparative scale reversed-phase high-pressure liquid chromatography (RP-HPLC) was performed on Agilent 1260 Infinity instruments equipped with a Waters xBridge Peptide BEH C18 column (5 μM particle size, 10 x 250 mm dimension) at a flow rate of 4 ml $min^{-1}$. Analytical scale RP-HPLC was performed on a Vydac 218ms C18 column (5 μM particle size, 4.6 x 150 mm dimension) at a flow rate of 1 ml $min^{-1}$. The mobile phase comprised 0.1% v/v trifluoroacetic acid in water (solvent A) and 90% acetonitrile, 0.1% TFA in water (solvent B).

Purification of $Cfa^C$-biotin and $Cfa^C$-Ir were performed using a 30-70% solvent B gradient. Pure fractions were identified by analytical RP-HPLC (0-70% solvent B gradient) and ESI-MS and were pooled and lyophilized. HPLC traces and ESI-MS spectra are provided in SI Figs. 1 & 2.

### Cloning

Plasmids of the general form **POI**-HA-Cfa$^{N}$-FLAG were cloned into a pcDNA3.1 vector backbone using Gibson Assembly (New England Biolabs) following the manufacturer's instructions. The encoded constructs were under the control of a CMV promoter. Point mutations were installed using QuikChange protocols (Agilent), following the manufacturer's instructions. Plasmid sequences were verified by Sanger sequencing (Genewiz) using universal primers. DNA sequences and corresponding amino acid sequences for the encoded proteins are provided below for each construct.

##### **H3.1**-HA-CFA$^{N}$-FLAG

```
ATGGCTCGTACCAAACAAACCGCGCGTAAGTCCACCGGCGGTAAAGCGCCACGTAAACAGCTGGCGACCAAAGCGGC
ACGCAAATCTGCGCCTGCGACCGGTGGTGTGAAAAAACCGCACCGTTACCGTCCGGGTACCGTTGCGCTGCGTGAGA
TCCGTCGTTACCAGAAGTCTACCGAACTGCTGATCCGTAAACTGCCGTTCCAGCGTCTGGTACGTGAAATCGCGCAG
GACTTCAAGACGGACCTGCGTTTCCAGTCTTCTGCGGTTATGGCGCTGCAAGAAGCGGCGGAAGCGTACCTGGTTGG
TCTGTTCGAAGATACCAACCTGGCGGCCATCCACGCTAAACGTGTTACCATCATGCCGAAAGACATCCAACTGGCGC
GTCGTATCCGTGGTGAACGTGCGGGTGGTTATCCGTATGATGTGCCGGATTATGCGTGCCTGTCTTACGACACAGAG
ATTCTGACCGTTGAATATGGATTCCTTCCTATCGGTAAGATCGTGGAGGAACGGATTGAATGCACAGTCTATACGGT
AGATAAAAATGGCTTTGTGTATACACAACCTATTGCTCAGTGGCATAACCGGGGAGAACAGGAAGTTTTCGAATACT
GCTTAGAAGACGGTTCGATTATCCGTGCAACGAAAGATCACAAATTTATGACGACCGACGGTCAGATGTTACCGATT
GATGAGATTTTCGAACGGGGGTTAGACCTGAAACAAGTTGATGGTTTGCCGAAAGGCGATTACAAGGATGACGACGA
TAAGTAA
```

```
MARTKQTARK STGGKAPRKQ LATKAARKSA PATGGVKKPH RYRPGTVALR EIRRYQKSTE
LLIRKLPFQR LVREIAQDFK TDLRFQSSAV MALQEAAEAY LVGLFEDTNL AAIHAKRVTI
MPKDIQLARR IRGERAGGYP YDVPDYACLS YDTEILTVEY GFLPIGKIVE ERIECTVYTV
DKNGFVYTQP IAQWHNRGEQ EVFEYCLEDG SIIRATKDHK FMTTDGQMLP IDEIFERGLD
LKQVDGLPKG DYKDDDDK
```

##### **CENP-A**-HA-CFA$^{N}$-FLAG

```
ATGGGGCCACGCAGGCGCAGTCGTAAACCAGAGGCGCCGCGACGGCGTTCGCCCAGCCCGACACCGACGCCGGGACC
GTCACGCCGTGGCCCTTCCTTGGGCGCAAGCAGCCACCAACACAGTAGACGCCGCCAAGGCTGGTTAAAGGAGATTC
GGAAGCTCCAGAAATCGACCCATTTGTTAATCCGTAAACTCCCGTTCTCTCGTCTAGCCCGCGAAATTTGCGTCAAA
TTCACTCGCGGCGTAGATTTCAACTGGCAGGCACAGGCGTTACTTGCACTTCAGGAAGCTGCGGAAGCGTTTCTGGT
GCATCTGTTTGAAGATGCTTATCTGCTGACCCTGCATGCCGGGCGTGTGACCCTGTTTCCTAAAGACGTTCAGCTGG
CCCGCCGTATCCGTGGTCTGGAAGAAGGTCTGGGTGGTTATCCGTATGATGTGCCGGATTATGCGTGCCTGTCTTAC
GACACAGAGATTCTGACCGTTGAATATGGATTCCTTCCTATCGGTAAGATCGTGGAGGAACGGATTGAATGCACAGT
CTATACGGTAGATAAAAATGGCTTTGTGTATACACAACCTATTGCTCAGTGGCATAACCGGGGAGAACAGGAAGTTT
TCGAATACTGCTTAGAAGACGGTTCGATTATCCGTGCAACGAAAGATCACAAATTTATGACGACCGACGGTCAGATG
TTACCGATTGATGAGATTTTCGAACGGGGGTTAGACCTGAAACAAGTTGATGGTTTGCCGAAAGGCGATTACAAGGA
TGACGACGATAAGTAA
```

```
MGPRRRSRKP EAPRRRSPSP TPTPGPSRRG PSLGASSHQH SRRRQGWLKE IRKLQKSTHL
LIRKLPFSRL AREICVKFTR GVDFNWQAQA LLALQEAAEA FLVHLFEDAY LLTLHAGRVT
LFPKDVQLAR RIRGLEEGLG GYPYDVPDYA CLSYDTEILT VEYGFLPIGK IVEERIECTV
YTVDKNGFVY TQPIAQWHNR GEQEVFEYCL EDGSIIRATK DHKFMTTDGQ MLPIDEIFER
GLDLKQVDGL PKGDYKDDDD K
```

#### **H4**-HA-CFA$^N$-FLAG

ATGTCTGGTCGTGGTAAAGGTGGTAAAGGTCTGGGTAAAGGTGGTGCTAAACGTCACCGTAAAGTTCTGCGTGACAA
CATCCAGGGTATCACCAAGCCGGCTATCCGTCGTCTGGCTCGTCGTGGTGGTGTTAAACGTATCTCCGGTCTGATCT
ACGAAGAAACCCGCGGTGTTCTGAAAGTTTTCCTGGAAAACGTTATCCGTGACGCTGTTACCTACACCGAACACGCT
AAACGTAAAACCGTTACCGCTATGGACGTTGTTTACGCTCTGAAACGTCAGGGTCGTACCCTGTACGGTTTCGGTGG
TTATCCGTATGATGTGCCGGATTATGCGTGCCTGTCTTACGACACAGAGATTCTGACCGTTGAATATGGATTCCTTC
CTATCGGTAAGATCGTGGAGGAACGGATTGAATGCACAGTCTATACGGTAGATAAAAATGGCTTTGTGTATACACAA
CCTATTGCTCAGTGGCATAACCGGGGAGAACAGGAAGTTTTCGAATACTGCTTAGAAGACGGTTCGATTATCCGTGC
AACGAAAGATCACAAATTTATGACGACCGACGGTCAGATGTTACCGATTGATGAGATTTTCGAACGGGGGTTAGACC
TGAAACAAGTTGATGGTTTGCCGAAAGGCGATTACAAGGATGACGACGATAAGTAA

MSGRGKGGKG LGKGGAKRHR KVLRDNIQGI TKPAIRRLAR RGGVKRISGL IYEETRGVLK
VFLENVIRDA VTYTEHAKRK TVTAMDVVYA LKRQGRTLYG FGGYPYDVPD YACLSYDTEI
LTVEYGFLPI GKIVEERIEC TVYTVDKNGF VYTQPIAQWH NRGEQEVFEY CLEDGSIIRA
TKDHKFMTTD GQMLPIDEIF ERGLDLKQVD GLPKGDYKDD DDK

#### **H1.1**-HA-CFA$^N$-FLAG

ATGTCTGAAACAGTGCCTCCCGCCCCCGCCGCTTCTGCTGCTCCTGAGAAACCTTTAGCTGGCAAGAAGGCAAAGAA
ACCTGCTAAGGCTGCAGCAGCCTCCAAGAAAAAACCCGCTGGCCCTTCCGTGTCAGAGCTGATCGTGCAGGCTGCTT
CCTCCTCTAAGGAGCGTGGTGGTGTGTCGTTGGCAGCTCTTAAAAAGGCGCTGGCGGCCGCAGGCTACGACGTGGAG
AAGAACAACAGCCGCATTAAGCTGGGCATTAAGAGCCTGGTAAGCAAGGGAACGTTGGTGCAGACAAAGGGTACCGG
AGCCTCGGGTTCCTTCAAGCTCAACAAGAAGGCGTCCTCCGTGGAAACCAAGCCCGGCGCCTCAAAGGTGGCTACAA
AAACTAAGGCAACGGGTGCATCTAAAAAGCTCAAAAAGGCCACGGGGGCTAGCAAAAAGAGCGTCAAGACTCCGAAA
AAGGCTAAAAAGCCTGCGGCAACAAGGAAATCCTCCAAGAATCCAAAAAAACCCAAAACTGTAAAGCCCAAGAAAGT
AGCTAAAAGCCCTGCTAAAGCTAAGGCTGTAAAACCCAAGGCGGCCAAGGCTAGGGTGACGAAGCCAAAGACTGCCA
AACCCAAGAAAGCGGCACCCAAGAAAAAGGGTGGTTATCCGTATGATGTGCCGGATTATGCGTGCCTGTCTTACGAC
ACAGAGATTCTGACCGTTGAATATGGATTCCTTCCTATCGGTAAGATCGTGGAGGAACGGATTGAATGCACAGTCTA
TACGGTAGATAAAAATGGCTTTGTGTATACACAACCTATTGCTCAGTGGCATAACCGGGGAGAACAGGAAGTTTTCG
AATACTGCTTAGAAGACGGTTCGATTATCCGTGCAACGAAAGATCACAAATTTATGACGACCGACGGTCAGATGTTA
CCGATTGATGAGATTTTCGAACGGGGGTTAGACCTGAAACAAGTTGATGGTTTGCCGAAAGGCGATTACAAGGATGA
CGACGATAAGTAA

MSETVPPAPA ASAAPEKPLA GKKAKKPAKA AAASKKKPAG PSVSELIVQA ASSSKERGGV
SLAALKKALA AAGYDVEKNN SRIKLGIKSL VSKGTLVQTK GTGASGSFKL NKKASSVETK
PGASKVATKT KATGASKKLK KATGASKKSV KTPKKAKKPA ATRKSSKNPK KPKTVKPKKV
AKSPAKAKAV KPKAAKARVT KPKTAKPKKA APKKKGGYPY DVPDYACLSY DTEILTVEYG
FLPIGKIVEE RIECTVYTVD KNGFVYTQPI AQWHNRGEQE VFEYCLEDGS IIRATKDHKF
MTTDGQMLPI DEIFERGLDL KQVDGLPKGD YKDDDDK

#### **H1.3**-HA-CFA$^N$-FLAG

ATGTCGGAGACTGCTCCACTTGCTCCTACCATTCCTGCACCCGCAGAAAAAACACCTGTGAAGAAAAAGGCGAAGAA
GGCAGGCGCAACTGCTGGGAAACGCAAAGCATCCGGACCCCCAGTATCTGAGCTTATCACCAAGGCAGTGGCAGCTT
CTAAGGAGCGCAGCGGCGTTTCTCTGGCCGCGCTTAAGAAAGCGCTTGCGGCTGCTGGCTACGATGTAGAAAAAAAC
AACAGCCGTATCAAGCTTGGCCTCAAGAGCTTGGTGAGCAAAGGTACTCTGGTGCAGACCAAAGGTACCGGTGCTTC
TGGCTCCTTCAAACTCAACAAGAAAGCGGCTTCCGGGGAAGGCAAACCCAAGGCCAAAAAGGCTGGCGCAGCCAAGC
CTAGGAAGCCTGCTGGGGCAGCCAAGAAGCCCAAGAAGGTGGCTGGCGCCGCTACCCCGAAGAAAAGCATCAAAAAG
ACTCCTAAGAAGGTAAAGAAGCCAGCAACCGCTGCTGGGACCAAGAAAGTGGCCAAGAGTGCGAAAAAGGTGAAAAC
ACCTCAGCCAAAAAAAGCTGCCAAGAGTCCAGCTAAGGCCAAAGCCCCTAAGCCCAAGGCGGCCAAGCCTAAGTCGG
GGAAGCCGAAGGTTACAAAGGCAAAGAAGGCAGCTCCGAAGAAAAAGGGTGGTTATCCGTATGATGTGCCGGATTAT
GCGTGCCTGTCTTACGACACAGAGATTCTGACCGTTGAATATGGATTCCTTCCTATCGGTAAGATCGTGGAGGAACG
GATTGAATGCACAGTCTATACGGTAGATAAAAATGGCTTTGTGTATACACAACCTATTGCTCAGTGGCATAACCGGG

GAGAACAGGAAGTTTTCGAATACTGCTTAGAAGACGGTTCGATTATCCGTGCAACGAAAGATCACAAATTTATGACG
ACCGACGGTCAGATGTTACCGATTGATGAGATTTTCGAACGGGGGTTAGACCTGAAACAAGTTGATGGTTTGCCGAA
AGGCGATTACAAGGATGACGACGATAAGTAA

MSETAPLAPT IPAPAEKTPV KKKAKKAGAT AGKRKASGPP VSELITKAVA ASKERSGVSL
AALKKALAAA GYDVEKNNSR IKLGLKSLVS KGTLVQTKGT GASGSFKLNK KAASGEGKPK
AKKAGAAKPR KPAGAAKKPK KVAGAATPKK SIKKTPKKVK KPATAAGTKK VAKSAKKVKT
PQPKKAAKSP AKAKAPKPKA AKPKSGKPKV TKAKKAAPKK KGGYPYDVPD YACLSYDTEI
LTVEYGFLPI GKIVEERIEC TVYTVDKNGF VYTQPIAQWH NRGEQEVFEY CLEDGSIIRA
TKDHKFMTTD GQMLPIDEIF ERGLDLKQVD GLPKGDYKDD DDK

**H2A**-HA-CFA$^{N}$-FLAG

ATGTCTGGACGTGGAAAGCAGGGAGGCAAGGCCCGCGCCAAGGCCAAGTCGCGCTCGTCCCGCGCTGGCCTTCAGTT
CCCGGTAGGGCGAGTGCATCGCTTGCTGCGCAAAGGCAACTACGCGGAGCGAGTGGGGGCCGGCGCGCCCGTCTACA
TGGCTGCAGTCCTCGAGTATCTGACCGCTGAGATCCTGGAGCTGGCGGGCAACGCGGCTCGGGACAACAAGAAGACG
CGCATCATCCCTCGTCACCTCCAGCTGGCCATCCGCAACGACGAGGAACTGAACAAGCTGCTGGGCAAAGTCACCAT
CGCCCAGGGCGGCGTCTTGCCTAACATCCAGGCCGTACTGCTCCCTAAGAAGACGGAGAGTCACCACAAGGCCAAGG
GCAAGGGTGGTTATCCGTATGATGTGCCGGATTATGCGTGCCTGTCTTACGACACAGAGATTCTGACCGTTGAATAT
GGATTCCTTCCTATCGGTAAGATCGTGGAGGAACGGATTGAATGCACAGTCTATACGGTAGATAAAAATGGCTTTGT
GTATACACAACCTATTGCTCAGTGGCATAACCGGGGAGAACAGGAAGTTTTCGAATACTGCTTAGAAGACGGTTCGA
TTATCCGTGCAACGAAAGATCACAAATTTATGACGACCGACGGTCAGATGTTACCGATTGATGAGATTTTCGAACGG
GGGTTAGACCTGAAACAAGTTGATGGTTTGCCGAAAGGCGATTACAAGGATGACGACGATAAGTAA

MSGRGKQGGK ARAKAKSRSS RAGLQFPVGR VHRLLRKGNY AERVGAGAPV YMAAVLEYLT
AEILELAGNA ARDNKKTRII PRHLQLAIRN DEELNKLLGK VTIAQGGVLP NIQAVLLPKK
TESHHKAKGK GGYPYDVPDY ACLSYDTEIL TVEYGFLPIG KIVEERIECT VYTVDKNGFV
YTQPIAQWHN RGEQEVFEYC LEDGSIIRAT KDHKFMTTDG QMLPIDEIFE RGLDLKQVDG
LPKGDYKDDD DK

**H2A-E92K**-HA-CFA$^{N}$-FLAG

ATGTCTGGACGTGGAAAGCAGGGAGGCAAGGCCCGCGCCAAGGCCAAGTCGCGCTCGTCCCGCGCTGGCCTTCAGTT
CCCGGTAGGGCGAGTGCATCGCTTGCTGCGCAAAGGCAACTACGCGGAGCGAGTGGGGGCCGGCGCGCCCGTCTACA
TGGCTGCAGTCCTCGAGTATCTGACCGCTGAGATCCTGGAGCTGGCGGGCAACGCGGCTCGGGACAACAAGAAGACG
CGCATCATCCCTCGTCACCTCCAGCTGGCCATCCGCAACGACGAGAAACTGAACAAGCTGCTGGGCAAAGTCACCAT
CGCCCAGGGCGGCGTCTTGCCTAACATCCAGGCCGTACTGCTCCCTAAGAAGACGGAGAGTCACCACAAGGCCAAGG
GCAAGGGTGGTTATCCGTATGATGTGCCGGATTATGCGTGCCTGTCTTACGACACAGAGATTCTGACCGTTGAATAT
GGATTCCTTCCTATCGGTAAGATCGTGGAGGAACGGATTGAATGCACAGTCTATACGGTAGATAAAAATGGCTTTGT
GTATACACAACCTATTGCTCAGTGGCATAACCGGGGAGAACAGGAAGTTTTCGAATACTGCTTAGAAGACGGTTCGA
TTATCCGTGCAACGAAAGATCACAAATTTATGACGACCGACGGTCAGATGTTACCGATTGATGAGATTTTCGAACGG
GGGTTAGACCTGAAACAAGTTGATGGTTTGCCGAAAGGCGATTACAAGGATGACGACGATAAGTAA

MSGRGKQGGK ARAKAKSRSS RAGLQFPVGR VHRLLRKGNY AERVGAGAPV YMAAVLEYLT
AEILELAGNA ARDNKKTRII PRHLQLAIRN DEKLNKLLGK VTIAQGGVLP NIQAVLLPKK
TESHHKAKGK GGYPYDVPDY ACLSYDTEIL TVEYGFLPIG KIVEERIECT VYTVDKNGFV
YTQPIAQWHN RGEQEVFEYC LEDGSIIRAT KDHKFMTTDG QMLPIDEIFE RGLDLKQVDG
LPKGDYKDDD DK

**SMC1A**-HA-CFA$^{N}$-FLAG

ATGGGGTTCCTGAAACTGATTGAGATTGAGAACTTTAAGTCGTACAAGGGTCGACAGATTATCGGACCATTTCAGAG
GTTCACCGCCATCATTGGACCCAATGGCTCTGGTAAGTCAAATCTCATGGATGCCATCAGCTTTGTGCTAGGTGAAA
AAACCAGCAACCTGCGGGTAAAGACCCTGCGGGACCTGATCCATGGAGCTCCTGTGGGCAAGCCAGCTGCCAACCGG
GCCTTTGTCAGCATGGTCTACTCTGAGGAGGGTGCTGAGGACCGTACCTTTGCCCGTGTCATTGTAGGAGGTTCTTC
TGAGTACAAGATCAACAACAAAGTGGTCCAACTACATGAGTACAGTGAGGAATTAGAGAAGTTGGGCATTCTCATCA

AAGCTCGTAACTTCCTCGTTTTCCAGGGTGCTGTGGAATCTATTGCCATGAAGAACCCCAAAGAGAGGACAGCTCTA
TTTGAAGAGATTAGTCGTTCTGGGGAGCTGGCGCAGGAGTATGACAAGCGAAAGAAGGAAATGGTGAAGGCTGAAGA
GGACACACAGTTTAATTACCATCGCAAGAAAAATATTGCGGCTGAACGCAAGGAAGCAAAGCAGGAGAAAGAAGAGG
CTGACCGGTACCAGCGCCTGAAGGATGAGGTAGTACGGGCTCAGGTACAGCTGCAGCTCTTTAAGCTTTACCATAAT
GAAGTGGAAATTGAGAAGCTCAACAAGGAACTGGCCTCAAAGAACAAGGAGATCGAGAAGGACAAGAAGCGTATGGA
CAAGGTGGAGGATGAACTGAAGGAGAAGAAGAAGGAGCTGGGCAAAATGATGCGGGAGCAGCAGCAGATTGAGAAGG
AGATCAAGGAGAAGGACTCAGAATTGAACCAGAAGCGGCCTCAGTACATCAAAGCCAAGGAGAACACCTCCCACAAA
ATCAAGAAGCTGGAAGCAGCCAAGAAGTCTCTGCAGAATGCTCAGAAGCACTACAAGAAGCGTAAAGGTGACATGGA
TGAGCTGGAGAAGGAGATGCTGTCAGTGGAGAAGGCTCGGCAGGAGTTTGAAGAACGGATGGAAGAAGAGAGTCAGA
GTCAGGGCAGAGATTTGACGTTGGAGGAGAATCAGGTGAAGAAATACCACCGGTTGAAAGAAGAAGCCAGCAAGAGA
GCAGCTACCCTGGCCCAGGAGCTGGAGAAATTCAATCGAGACCAGAAAGCTGACCAGGACCGTCTGGATCTGGAAGA
ACGGAAGAAAGTAGAGACAGAGGCCAAGATCAAGCAAAAGCTGCGGGAAATTGAAGAGAATCAGAAGCGGATTGAGA
AACTGGAGGAATACATCACCACTAGCAAGCAGTCCCTAGAAGAGCAGAAGAAGCTAGAGGGGGAGCTGACAGAGGAG
GTGGAGATGGCCAAGCGGCGTATTGATGAAATCAATAAGGAGCTGAACCAGGTGATGGAGCAGCTAGGGGATGCCCG
CATCGACCGCCAGGAGAGCAGCCGCCAGCAGCGAAAGGCAGAGATAATGGAAAGCATCAAGCGCCTTTACCCTGGCT
CTGTGTACGGCCGCCTCATTGACCTATGCCAGCCCACACAAAAGAAGTATCAGATTGCTGTAACCAAGGTTTTGGGC
AAGAACATGGATGCCATTATTGTGGACTCGGAGAAGACAGGCCGGGACTGTATTCAGTATATCAAGGAGCAGCGTGG
GGAGCCTGAGACCTTCTTGCCTCTTGACTACCTGGAGGTGAAGCCTACAGATGAGAAACTCCGGGAGCTGAAGGGGG
CCAAGCTAGTGATTGATGTGATTCGCTATGAGCCACCTCATATCAAAAAGGCCCTGCAGTATGCTTGTGGCAATGCC
CTTGTCTGTGACAACGTGGAAGATGCCCGCCGCATTGCCTTTGGAGGCCACCAGCGCCACAAGACAGTGGCACTGGA
TGGAACCCTATTCCAGAAGTCAGGAGTGATCTCTGGTGGGGCCAGTGACCTGAAGGCCAAGGCACGGCGCTGGGATG
AGAAAGCAGTAGACAAGTTGAAAGAGAAGAAGGAGCGCTTGACAGAGGAGCTGAAAGAGCAGATGAAGGCAAAACGG
AAAGAGGCAGAGCTGCGTCAGGTGCAGTCTCAGGCCCATGGACTGCAGATGCGGCTCAAGTACTCCCAGAGTGACCT
AGAACAGACCAAGACACGACATCTAGCCCTGAATCTGCAGGAAAAATCCAAGCTGGAGAGTGAGCTAGCCAACTTTG
GGCCTCGCATTAATGATATCAAGAGGATCATTCAGAGCCGAGAGAGGGAAATGAAAGACTTGAAGGAGAAGATGAAC
CAGGTAGAGGATGAGGTGTTTGAAGAGTTTTGTCGGGAGATTGGTGTGCGCAACATCCGGGAGTTTGAGGAAGAAAA
GGTGAAACGGCAGAATGAAATCGCCAAGAAGCGTTTGGAGTTTGAGAATCAGAAGACTCGCTTGGGCATTCAGTTGG
ATTTTGAAAAGAACCAACTGAAGGAGGACCAAGATAAAGTACACATGTGGGAGCAGACAGTGAAAAAAGATGAAAAT
GAGATAGAAAAGCTCAAAAAGGAGGAACAAAGACACATGAAGATCATAGATGAGACCATGGCTCAGCTACAAGACCT
GAAGAATCAGCATCTGGCCAAGAAGTCGGAAGTGAATGACAAGAATCATGAGATGGAGGAGATTCGTAAGAAACTCG
GGGGCGCCAACAAGGAAATGACCCATTTACAGAAGGAGGTGACAGCCATTGAGACCAAGCTTGAACAGAAGCGCAGT
GACCGTCACAACTTGCTACAGGCCTGTAAGATGCAGGACATTAAGTTGCCACTGTCAAAAGGCACCATGGATGATAT
TAGTCAGGAAGAGGGTAGCTCCCAGGGGGGAGGACTCAGTGAGTGGTTCACAGAGAATTTCCAGTATCTATGCACGAG
AGGCCCTCATTGAGATTGACTACGGTGATCTGTGTGAGGATCTGAAGGATGCCCAGGCTGAGGAAGAGATCAAGCAA
GAGATGAACACACTGCAGCAGAAGCTGAATGAGCAGCAGAGTGTGCTTCAGCGTATTGCCGCCCCCAACATGAAGGC
CATGGAAAAGCTGGAAAGTGTCCGAGACAAGTTCCAGGAGACCTCAGATGAGTTTGAAGCAGCCCGAAAGCGAGCAA
AGAAGGCCAAGCAGGCATTCGAACAGATCAAGAAGGAGCGCTTTGACCGCTTCAATGCTTGTTTTGAATCTGTGGCT
ACCAACATTGATGAGATCTATAAGGCCCTGTCCCGCAATAGCAGTGCCCAGGCATTCCTGGGCCCTGAGAACCCTGA
AGAGCCCTACTTGGATGGCATCAACTACAACTGTGTGGCTCCTGGGAAACGCTTCCGGCCTATGGACAACTTGTCAG
GCGGGGAGAAGACAGTGGCAGCTCTGGCCCTGCTCTTTGCCATCCACAGCTACAAGCCAGCCCCCTTCTTCGTCCTG
GATGAGATTGATGCTGCCTTGGATAACACCAACATTGGCAAGGTGGCAAATTACATCAAGGAGCAGTCGACTTGCAA
CTTCCAGGCCATCGTCATCTCTCTCAAGGAGGAGTTCTACACCAAGGCCGAGAGCCTCATTGGAGTCTATCCTGAGC
AAGGGGACTGTGTGATCAGCAAAGTCCTGACCTTCGACCTCACCAAGTACCCAGATGCCAACCCCAACCCCAATGAG
CAGGGTGGTTATCCGTATGATGTGCCGGATTATGCGTGCCTGTCTTACGACACAGAGATTCTGACCGTTGAATATGG
ATTCCTTCCTATCGGTAAGATCGTGGAGGAACGGATTGAATGCACAGTCTATACGGTAGATAAAAATGGCTTTGTGT
ATACACAACCTATTGCTCAGTGGCATAACCGGGGAGAACAGGAAGTTTTCGAATACTGCTTAGAAGACGGTTCGATT
ATCCGTGCAACGAAAGATCACAAATTTATGACGACCGACGGTCAGATGTTACCGATTGATGAGATTTTCGAACGGGG
GTTAGACCTGAAACAAGTTGATGGTTTGCCGAAAGGCGATTACAAGGATGACGACGATAAGTAA

MGFLKLIEIE NFKSYKGRQI IGPFQRFTAI IGPNGSGKSN LMDAISFVLG EKTSNLRVKT
LRDLIHGAPV GKPAANRAFV SMVYSEEGAE DRTFARVIVG GSSEYKINNK VVQLHEYSEE
LEKLGILIKA RNFLVFQGAV ESIAMKNPKE RTALFEEISR SGELAQEYDK RKKEMVKAEE
DTQFNYHRKK NIAAERKEAK QEKEEADRYQ RLKDEVVRAQ VQLQLFKLYH NEVEIEKLNK
ELASKNKEIE KDKKRMDKVE DELKEKKKEL GKMMREQQQI EKEIKEKDSE LNQKRPQYIK
AKENTSHKIK KLEAAKKSLQ NAQKHYKKRK GDMDELEKEM LSVEKARQEF EERMEEESQS
QGRDLTLEEN QVKKYHRLKE EASKRAATLA QELEKFNRDQ KADQDRLDLE ERKKVETEAK

IKQKLREIEE NQKRIEKLEE YITTSKQSLE EQKKLEGELT EEVEMAKRRI DEINKELNQV
MEQLGDARID RQESSRQQRK AEIMESIKRL YPGSVYGRLI DLCQPTQKKY QIAVTKVLGK
NMDAIIVDSE KTGRDCIQYI KEQRGEPETF LPLDYLEVKP TDEKLRELKG AKLVIDVIRY
EPPHIKKALQ YACGNALVCD NVEDARRIAF GGHQRHKTVA LDGTLFQKSG VISGGASDLK
AKARRWDEKA VDKLKEKKER LTEELKEQMK AKRKEAELRQ VQSQAHGLQM RLKYSQSDLE
QTKTRHLALN LQEKSKLESE LANFGPRIND IKRIIQSRER EMKDLKEKMN QVEDEVFEEF
CREIGVRNIR EFEEEKVKRQ NEIAKKRLEF ENQKTRLGIQ LDFEKNQLKE DQDKVHMWEQ
TVKKDENEIE KLKKEEQRHM KIIDETMAQL QDLKNQHLAK KSEVNDKNHE MEEIRKKLGG
ANKEMTHLQK EVTAIETKLE QKRSDRHNLL QACKMQDIKL PLSKGTMDDI SQEEGSSQGE
DSVSGSQRIS SIYAREALIE IDYGDLCEDL KDAQAEEEIK QEMNTLQQKL NEQQSVLQRI
AAPNMKAMEK LESVRDKFQE TSDEFEAARK RAKKAKQAFE QIKKERFDRF NACFESVATN
IDEIYKALSR NSSAQAFLGP ENPEEPYLDG INYNCVAPGK RFRPMDNLSG GEKTVAALAL
LFAIHSYKPA PFFVLDEIDA ALDNTNIGKV ANYIKEQSTC NFQAIVISLK EEFYTKAESL
IGVYPEQGDC VISKVLTFDL TKYPDANPNP NEQGGYPYDV PDYACLSYDT EILTVEYGFL
PIGKIVEERI ECTVYTVDKN GFVYTQPIAQ WHNRGEQEVF EYCLEDGSII RATKDHKFMT
TDGQMLPIDE IFERGLDLKQ VDGLPKGDYK DDDDK

**CTCF**-HA-CFA$^N$-FLAG

ATGGAAGGTGATGCAGTCGAAGCCATTGTGGAGGAGTCCGAAACTTTTATTAAAGGAAAGGAGAGAAAGACTTACCA
GAGACGCCGGGAAGGGGGCCAGGAAGAAGATGCCTGCCACTTACCCCAGAACCAGACGGATGGGGGTGAGGTGGTCC
AGGATGTCAACAGCAGTGTACAGATGGTGATGATGGAACAGCTGGACCCCACCCTTCTTCAGATGAAGACTGAAGTA
ATGGAGGGCACAGTGGCTCCAGAAGCAGAGGCTGCTGTGGACGATACCCAGATTATAACTTTACAGGTTGTAAATAT
GGAGGAACAGCCCATAAACATAGGAGAACTTCAGCTTGTTCAAGTACCTGTTCCTGTGACTGTACCTGTTGCTACCA
CTTCAGTAGAAGAACTTCAGGGGGCTTATGAAAATGAAGTGTCTAAAGAGGGCCTTGCGGAAAGTGAACCCATGATA
TGCCACACCCTACCTTTGCCTGAAGGGTTTCAGGTGGTTAAAGTGGGGGCCAATGGAGAGGTGGAGACACTAGAACA
AGGGGAACTTCCACCCCAGGAAGATCCTAGTTGGCAAAAAGACCCAGACTATCAGCCACCAGCCAAAAAAACAAAGA
AAACCAAAAAGAGCAAACTGCGTTATACAGAGGAGGGCAAAGATGTAGATGTGTCTGTCTACGATTTTGAGGAAGAA
CAGCAGGAGGGTCTGCTATCAGAGGTTAATGCAGAGAAAGTGGTTGGTAATATGAAGCCTCCAAAGCCAACAAAAAT
TAAAAAGAAAGGTGTAAAGAAGACATTCCAGTGTGAGCTTTGCAGTTACACGTGTCCACGGCGTTCAAATTTGGATC
GTCACATGAAAAGCCACACTGATGAGAGACCACACAAGTGCCATCTCTGTGGCAGGGCATTCAGAACAGTCACCCTC
CTGAGGAATCACCTTAACACACACACAGGTACTCGTCCTCACAAGTGCCCAGACTGCGACATGGCCTTTGTGACCAG
TGGAGAATTGGTTCGGCATCGTCGTTACAAACACACCCACGAGAAGCCATTCAAGTGTTCCATGTGCGATTACGCCA
GTGTAGAAGTCAGCAAATTAAAACGTCACATTCGCTCTCATACTGGAGAGCGTCCGTTTCAGTGCAGTTTGTGCAGT
TATGCCAGCAGGGACACATACAAGCTGAAAAGGCACATGAGAACCCATTCAGGGGAAAAGCCTTATGAATGTTATAT
TTGTCATGCTCGGTTTACCCAAAGTGGTACCATGAAGATGCACATTTTACAGAAGCACACAGAAAATGTGGCCAAAT
TTCACTGTCCCCACTGTGACACAGTCATAGCCCGAAAAAGTGATTTGGGTGTCCACTTGCGAAAGCAGCATTCCTAT
ATTGAGCAAGGCAAGAAATGCCGTTACTGTGATGCTGTGTTTCATGAGCGCTATGCCCTCATCCAGCATCAGAAGTC
ACACAAGAATGAGAAGCGCTTTAAGTGTGACCAGTGTGATTACGCTTGTAGACAGGAGAGGCACATGATCATGCACA
AGCGCACCCACACCGGGGAGAAGCCTTACGCCTGCAGCCACTGCGATAAGACCTTCCGCCAGAAGCAGCTTCTCGAC
ATGCACTTCAAGCGCTATCACGACCCCAACTTCGTCCCTGCGGCTTTTGTCTGTTCTAAGTGTGGGAAAACATTTAC
ACGTCGGAATACCATGGCAAGACATGCTGATAATTGTGCTGGCCCAGATGGCGTAGAGGGGGAAAATGGAGGAGAAA
CGAAGAAGAGTAAACGTGGAAGAAAAAGAAAGATGCGCTCTAAGAAAGAAGATTCCTCTGACAGTGAAAATGCTGAA
CCAGATCTGGACGACAATGAGGATGAGGAGGAGCCTGCCGTAGAAATTGAACCTGAGCCAGAGCCTCAGCCTGTGAC
CCCAGCCCCACCACCCGCCAAGAAGCGGAGAGGACGACCCCCTGGCAGAACCAACCAGCCCAAACAGAACCAGCCAA
CAGCTATCATTCAGGTTGAAGACCAGAATACAGGTGCAATTGAGAACATTATAGTTGAAGTAAAAAAAGAGCCAGAT
GCTGAGCCCGCAGAGGGAGAGGAAGAGGAGGCCCAGCCAGCTGCCACAGATGCCCCCAACGGAGACCTCACGCCCGA
GATGATCCTCAGCATGATGGACCGGGGTGGTTATCCGTATGATGTGCCGGATTATGCGTGCCTGTCTTACGACACAG
AGATTCTGACCGTTGAATATGGATTCCTTCCTATCGGTAAGATCGTGGAGGAACGGATTGAATGCACAGTCTATACG
GTAGATAAAAATGGCTTTGTGTATACACAACCTATTGCTCAGTGGCATAACCGGGGAGAACAGGAAGTTTTCGAATA
CTGCTTAGAAGACGGTTCGATTATCCGTGCAACGAAAGATCACAAATTTATGACGACCGACGGTCAGATGTTACCGA
TTGATGAGATTTTCGAACGGGGGTTAGACCTGAAACAAGTTGATGGTTTGCCGAAAGGCGATTACAAGGATGACGAC
GATAAGTAA

MEGDAVEAIV EESETFIKGK ERKTYQRRRE GGQEEDACHL PQNQTDGGEV VQDVNSSVQM
VMMEQLDPTL LQMKTEVMEG TVAPEAEAAV DDTQIITLQV VNMEEQPINI GELQLVQVPV

PVTVPVATTS VEELQGAYEN EVSKEGLAES EPMICHTLPL PEGFQVVKVG ANGEVETLEQ
GELPPQEDPS WQKDPDYQPP AKKTKKTKKS KLRYTEEGKD VDVSVYDFEE EQQEGLLSEV
NAEKVVGNMK PPKPTKIKKK GVKKTFQCEL CSYTCPRRSN LDRHMKSHTD ERPHKCHLCG
RAFRTVTLLR NHLNTHTGTR PHKCPDCDMA FVTSGELVRH RRYKHTHEKP FKCSMCDYAS
VEVSKLKRHI RSHTGERPFQ CSLCSYASRD TYKLKRHMRT HSGEKPYECY ICHARFTQSG
TMKMHILQKH TENVAKFHCP HCDTVIARKS DLGVHLRKQH SYIEQGKKCR YCDAVFHERY
ALIQHQKSHK NEKRFKCDQC DYACRQERHM IMHKRTHTGE KPYACSHCDK TFRQKQLLDM
HFKRYHDPNF VPAAFVCSKC GKTFTRRNTM ARHADNCAGP DGVEGENGGE TKKSKRGRKR
KMRSKKEDSS DSENAEPDLD DNEDEEEPAV EIEPEPEPQP VTPAPPPAKK RRGRPPGRTN
QPKQNQPTAI IQVEDQNTGA IENIIVEVKK EPDAEPAEGE EEEAQPAATD APNGDLTPEM
ILSMMDRGGY PYDVPDYACL SYDTEILTVE YGFLPIGKIV EERIECTVYT VDKNGFVYTQ
PIAQWHNRGE QEVFEYCLED GSIIRATKDH KFMTTDGQML PIDEIFERGL DLKQVDGLPK
GDYKDDDDK

**RPB1**-HA-CFA$^{N}$-FLAG

ATGCACGGGGGTGGCCCCCCCTCGGGGGACAGCGCATGCCCGCTGCGCACCATCAAGAGAGTCCAGTTCGGAGTCCT
GAGTCCGGATGAACTGAAGCGAATGTCTGTGACGGAGGGTGGCATCAAATACCCAGAGACGACTGAGGGAGGCCGCC
CCAAGCTTGGGGGGCTGATGGACCCGAGGCAGGGGGTGATTGAGCGGACTGGCCGCTGCCAAACATGTGCAGGAAAC
ATGACAGAGTGTCCTGGCCACTTTGGCCACATTGAACTGGCCAAGCCTGTGTTTCACGTGGGCTTCCTGGTGAAGAC
AATGAAAGTTTTGCGCTGTGTCTGCTTCTTCTGCTCCAAACTGCTTGTGGACTCTAACAACCCAAAGATCAAGGATA
TCCTGGCTAAGTCCAAGGGACAGCCCAAGAAGCGGCTCACACATGTCTACGACCTTTGCAAGGGCAAAAACATATGC
GAGGGTGGGGAGGAGATGGACAACAAGTTCGGTGTGGAACAACCTGAGGGTGACGAGGATCTGACCAAAGAAAAGGG
CCATGGTGGCTGTGGGCGGTACCAGCCCAGGATCCGGCGTTCTGGCCTAGAGCTGTATGCGGAATGGAAGCACGTTA
ATGAGGACTCTCAGGAGAAGAAGATCCTGCTGAGTCCAGAGCGAGTGCATGAGATCTTCAAACGCATCTCAGATGAG
GAGTGTTTTGTGCTGGGCATGGAGCCCCGCTATGCACGGCCAGAGTGGATGATTGTCACAGTGCTGCCTGTGCCCCC
GCTCTCCGTGCGGCCTGCTGTTGTGATGCAGGGCTCTGCCCGTAACCAGGATGACCTGACTCACAAACTGGCTGACA
TCGTGAAGATCAACAATCAGCTGCGGCGCAATGAGCAGAACGGCGCAGCGGCCCATGTCATTGCAGAGGATGTGAAG
CTCCTCCAGTTCCATGTGGCCACCATGGTGGACAATGAGCTGCCTGGCTTGCCCCGTGCCATGCAGAAGTCTGGGCG
TCCCCTCAAGTCCCTGAAGCAGCGGTTGAAGGGCAAGGAAGGCCGGGTGCGAGGGAACCTGATGGGCAAAAGAGTGG
ACTTCTCGGCCCGTACTGTCATCACCCCCGACCCCAACCTCTCCATTGACCAGGTTGGCGTGCCCCGCTCCATTGCT
GCCAACATGACCTTTGCGGAGATTGTCACCCCCTTCAACATTGACAGACTTCAAGAACTAGTGCGCAGGGGGAACAG
CCAGTACCCAGGCGCCAAGTACATCATCCGAGACAATGGTGATCGCATTGACTTGCGTTTCCACCCCAAGCCCAGTG
ACCTTCACCTGCAGACCGGCTATAAGGTGGAACGGCACATGTGTGATGGGGACATTGTTATCTTCAACCGGCAGCCA
ACTCTGCACAAAATGTCCATGATGGGGCATCGGGTCCGCATTCTCCCATGGTCTACCTTTCGCTTGAATCTTAGCGT
GACAACTCCGTACAATGCAGACTTTGACGGGGATGAGATGAACTTGCACCTGCCACAGTCTCTGGAGACGCGAGCAG
AGATCCAGGAGCTGGCCATGGTTCCTCGCATGATTGTCACCCCCCAGAGCAATCGGCCTGTCATGGGTATTGTGCAG
GACACACTCACAGCAGTGCGCAAATTCACCAAGAGAGACGTCTTCCTGGAGCGGGGTGAAGTGATGAACCTCCTGAT
GTTCCTGTCGACGTGGGATGGGAAGGTCCCACAGCCGGCCATCCTAAAGCCCCGGCCCCTGTGGACAGGCAAGCAAA
TCTTCTCCCTCATCATACCTGGTCACATCAATTGTATCCGTACCCACAGCACCCATCCCGATGATGAAGACAGTGGC
CCTTACAAGCACATCTCTCCTGGGGACACCAAGGTGGTGGTGGAGAATGGGGAGCTGATCATGGGCATCCTGTGTAA
GAAGTCTCTGGGCACGTCAGCTGGCTCCCTGGTCCACATCTCCTACCTAGAGATGGGTCATGACATCACTCGCCTCT
TCTACTCCAACATTCAGACTGTCATTAACAACTGGCTCCTCATCGAGGGTCATACTATTGGCATTGGGGACTCCATT
GCTGATTCTAAGACTTACCAGGACATTCAGAACACTATTAAGAAGGCCAAGCAGGACGTAATAGAGGTCATCGAGAA
GGCACACAACAATGAGCTGGAGCCCACCCCAGGGAACACTCTGCGGCAGACGTTTGAGAATCAGGTGAACCGCATTC
TTAACGATGCCCGAGACAAGACTGGCTCCTCTGCTCAGAAATCCCTGTCTGAATACAACAACTTCAAGTCTATGGTC
GTGTCCGGAGCTAAAGGTTCCAAGATTAACATCTCCCAGGTCATTGCTGTCGTTGGACAGCAGGACGTCGAGGGCAA
GCGGATTCCATTTGGCTTCAAGCACCGGACTCTGCCTCACTTCATCAAGGATGACTACGGGCCTGAGAGCCGTGGCT
TTGTGGAGAACTCCTACCTAGCCGGCCTCACACCCACTGAGTTCTTTTTCCACGCCATGGGGGGTCGTGAGGGGCTC
ATTGACACGGCTGTCAAGACTGCTGAGACTGGATACATCCAGCGGCGGCTGATCAAGTCCATGGAGTCAGTGATGGT
GAAGTACGACGCGACTGTGCGGAACTCCATCAACCAGGTGGTGCAGCTGCGCTACGGCGAAGACGGCCTGGCAGGCG
AGAGCGTTGAGTTCCAGAACCTGGCTACGCTTAAGCCTTCCAACAAGGCTTTTGAGAAGAAGTTCCGCTTTGATTAT
ACCAATGAGAGGGCCCTGCGGCGCACTCTGCAGGAGGACCTGGTGAAGGACGTGCTGAGCAACGCACACATCCAGAA
CGAGTTGGAGCGGGAATTTGAGCGGATGCGGGAGGATCGGGAGGTGCTCAGGGTCATCTTCCCAACTGGAGACAGCA
AGGTCGTCCTCCCCCTGTAACCTGCTGCGGATGATCTGGAATGCTCAGAAAATCTTCCACATCAACCCACGCCTTCCC
TCCGACCTGCACCCCATCAAAGTGGTGGAGGGAGTCAAGGAATTGAGCAAGAAGCTGGTGATTGTGAATGGGGATGA

CCCACTAAGTCGACAGGCCCAGGAAAATGCCACGCTGCTCTTCAACATCCACCTGCGGTCCACGTTGTGTTCCCGCC
GCATGGCAGAGGAGTTTCGGCTCAGTGGGGAGGCCTTCGACTGGCTGCTTGGGGAGATTGAGTCCAAGTTCAACCAA
GCCATTGCGCATCCCGGGGAAATGGTGGGGGCTCTGGCTGCGCAGTCCCTTGGAGAACCTGCCACCCAGATGACCTT
GAATACCTTCCACTATGCTGGTGTGTCTGCCAAGAATGTGACGCTGGGTGTGCCCCGACTTAAGGAGCTCATCAACA
TTTCCAAGAAGCCAAAGACTCCTTCGCTTACTGTCTTCCTGTTGGGCCAGTCCGCTCGAGATGCTGAGAGAGCCAAG
GATATTCTGTGCCGTCTGGAGCATACAACGTTGAGGAAGGTGACTGCCAACACAGCCATCTACTATGACCCCAACCC
CCAGAGCACGGTGGTGGCAGAGGATCAGGAATGGGTGAATGTCTACTATGAAATGCCTGACTTTGATGTGGCCCGAA
TCTCCCCCTGGCTGTTGCGGGTGGAGCTGGATCGGAAGCACATGACTGACCGGAAGCTCACCATGGAGCAGATTGCT
GAAAAGATCAATGCTGGTTTTGGTGACGACTTGAACTGCATCTTTAATGATGACAATGCAGAGAAGCTGGTGCTCCG
TATTCGCATCATGAACAGCGATGAGAACAAGATGCAAGAGGAGGAAGAGGTGGTGGACAAGATGGATGATGATGTCT
TCCTGCGCTGCATCGAGTCCAACATGCTGACAGATATGACCCTGCAGGGCATCGAGCAGATCAGCAAGGTGTACATG
CACTTGCCACAGACAGACAACAAGAAGAAGATCATCATCACGGAGGATGGGGAATTCAAGGCCCTGCAGGAGTGGAT
CCTGGAGACGGACGGCGTGAGCTTGATGCGGGTGCTGAGTGAGAAGGACGTGGACCCCGTACGCACCACGTCCAATG
ACATTGTGGAGATCTTCACGGTGCTGGGCATTGAAGCCGTGCGGAAGGCCCTGGAGCGGGAGCTGTACCACGTCATC
TCCTTTGATGGCTCCTATGTCAATTACCGACACTTGGCTCTCTTGTGTGATACCATGACCTGTCGTGGCCACTTGAT
GGCCATCACCCGACACGGAGTCAACCGCCAGGACACAGGACCACTCATGAAGTGTTCCTTTGAGGAAACGGTGGACG
TGCTTATGGAAGCAGCCGCACACGGTGAGAGTGACCCCATGAAGGGGGTCTCTGAGAATATCATGCTGGGCCAGCTG
GCTCCGGCCGGCACTGGCTGCTTTGACCTCCTGCTTGATGCAGAGAAGTGCAAGTATGGCATGGAGATCCCCACCAA
TATCCCCGGCCTGGGGGCTGCTGGACCCACCGGCATGTTCTTTGGTTCAGCACCCAGTCCCATGGGTGGAATCTCTC
CTGCCATGACACCTTGGAACCAGGGTGCAACCCCTGCCTATGGCGCCTGGTCCCCCAGTGTTGGGAGTGGAATGACC
CCAGGGGCAGCCGGCTTCTCTCCCAGTGCTGCGTCAGATGCCAGCGGCTTCAGCCCAGGTTACTCCCCTGCCTGGTC
TCCCACACCGGGCTCCCCGGGGTCCCCAGGTCCCTCAAGCCCCTACATCCCTTCACCAGGTGGTGCCATGTCTCCCA
GCTACTCGCCAACGTCACCTGCCTACGAGCCCCGCTCTCCTGGGGGCTACACACCCCAGAGTCCCTCTTATTCCCCC
ACTTCACCCTCCTACTCCCCTACCTCTCCATCCTATTCTCCAACCAGTCCCAACTATAGTCCCACATCACCCAGCTA
TTCGCCAACGTCACCCAGCTACTCACCGACCTCTCCCAGCTACTCACCCACCTCTCCCAGCTACTCGCCCACCTCTC
CCAGCTACTCGCCCACCTCTCCCAGCTACTCACCCACTTCCCCTAGCTACTCGCCCACTTCCCCTAGCTACTCGCCA
ACGTCTCCCAGCTACTCGCCGACATCTCCCAGCTACTCGCCAACTTCACCCAGCTATTCTCCCACTTCTCCCAGCTA
CTCACCTACCTCTCCAAGCTATTCACCCACCTCCCCCAGCTACTCACCCACTTCCCCAAGTTACTCACCCACCAGCC
CGAACTATTCTCCAACCAGTCCCAATTACACCCCAACATCACCCAGCTACAGCCCGACATCACCCAGCTATTCACCT
ACTAGTCCCAACTACACACCTACCAGCCCTAACTACAGCCCAACCTCTCCAAGCTACTCTCCAACATCACCCAGCTA
TTCCCCGACCTCACCAAGTTACTCCCCTTCCAGCCCACGATACACACCACAGTCTCCAACCTATACCCCAAGCTCAC
CCAGCTACAGCCCCAGCTCGCCCAGCTACAGCCCAACCTCACCCAAGTACACCCCAACCAGTCCTTCTTACAGTCCC
AGCTCCCCAGAGTATACCCCAACCTCTCCCAAGTACTCACCTACCAGTCCCAAATATTCACCCACCTCTCCCAAGTA
CTCGCCTACCAGTCCCACCTATTCACCCACCACCCCAAAATACTCCCCAACATCTCCTACTTATTCCCCAACCTCTC
CAGTCTACACCCCAACCTCTCCCAAGTACTCACCTACTAGCCCCACTTACTCGCCCACTTCCCCCAAGTACTCGCCC
ACCAGCCCCACCTACTCGCCCACCTCCCCCAAAGGCTCAACCTACTCTCCCACTTCCCCTGGTTACTCGCCCACCAG
CCCCACCTACAGTCTCACAAGCCCGGCTATCAGCCCGGATGACAGTGACGAGGAGAACGCTAGCGGTGGTTATCCGT
ATGATGTGCCGGATTATGCGTGCCTGTCTTACGACACAGAGATTCTGACCGTTGAATATGGATTCCTTCCTATCGGT
AAGATCGTGGAGGAACGGATTGAATGCACAGTCTATACGGTAGATAAAAATGGCTTTGTGTATACACAACCTATTGC
TCAGTGGCATAACCGGGGAGAACAGGAAGTTTTCGAATACTGCTTAGAAGACGGTTCGATTATCCGTGCAACGAAAG
ATCACAAATTTATGACGACCGACGGTCAGATGTTACCGATTGATGAGATTTTCGAACGGGGGTTAGACCTGAAACAA
GTTGATGGTTTGCCGAAAGGCGATTACAAGGATGACGACGATAAGTAA

MHGGGPPSGD SACPLRTIKR VQFGVLSPDE LKRMSVTEGG IKYPETTEGG RPKLGGLMDP
RQGVIERTGR CQTCAGNMTE CPGHFGHIEL AKPVFHVGFL VKTMKVLRCV CFFCSKLLVD
SNNPKIKDIL AKSKGQPKKR LTHVYDLCKG KNICEGGEEM DNKFGVEQPE GDEDLTKEKG
HGGCGRYQPR IRRSGLELYA EWKHVNEDSQ EKKILLSPER VHEIFKRISD EECFVLGMEP
RYARPEWMIV TVLPVPPLSV RPAVVMQGSA RNQDDLTHKL ADIVKINNQL RRNEQNGAAA
HVIAEDVKLL QFHVATMVDN ELPGLPRAMQ KSGRPLKSLK QRLKGKEGRV RGNLMGKRVD
FSARTVITPD PNLSIDQVGV PRSIAANMTF AEIVTPFNID RLQELVRRGN SQYPGAKYII
RDNGDRIDLR FHPKPSDLHL QTGYKVERHM CDGDIVIFNR QPTLHKMSMM GHRVRILPWS
TFRLNLSVTT PYNADFDGDE MNLHLPQSLE TRAEIQELAM VPRMIVTPQS NRPVMGIVQD
TLTAVRKFTK RDVFLERGEV MNLLMFLSTW DGKVPQPAIL KPRPLWTGKQ IFSLIIPGHI
NCIRTHSTHP DDEDSGPYKH ISPGDTKVVV ENGELIMGIL CKKSLGTSAG SLVHISYLEM
GHDITRLFYS NIQTVINNWL LIEGHTIGIG DSIADSKTYQ DIQNTIKKAK QDVIEVIEKA
HNNELEPTPG NTLRQTFENQ VNRILNDARD KTGSSAQKSL SEYNNFKSMV VSGAKGSKIN

```
ISQVIAVVGQ QDVEGKRIPF GFKHRTLPHF IKDDYGPESR GFVENSYLAG LTPTEFFFHA
MGGREGLIDT AVKTAETGYI QRRLIKSMES VMVKYDATVR NSINQVVQLR YGEDGLAGES
VEFQNLATLK PSNKAFEKKF RFDYTNERAL RRTLQEDLVK DVLSNAHIQN ELEREFERMR
EDREVLRVIF PTGDSKVVLP CNLLRMIWNA QKIFHINPRL PSDLHPIKVV EGVKELSKKL
VIVNGDDPLS RQAQENATLL FNIHLRSTLC SRRMAEEFRL SGEAFDWLLG EIESKFNQAI
AHPGEMVGAL AAQSLGEPAT QMTLNTFHYA GVSAKNVTLG VPRLKELINI SKKPKTPSLT
VFLLGQSARD AERAKDILCR LEHTTLRKVT ANTAIYYDPN PQSTVVAEDQ EWVNVYYEMP
DFDVARISPW LLRVELDRKH MTDRKLTMEQ IAEKINAGFG DDLNCIFNDD NAEKLVLRIR
IMNSDENKMQ EEEEVVDKMD DDVFLRCIES NMLTDMTLQG IEQISKVYMH LPQTDNKKKI
IITEDGEFKA LQEWILETDG VSLMRVLSEK DVDPVRTTSN DIVEIFTVLG IEAVRKALER
ELYHVISFDG SYVNYRHLAL LCDTMTCRGH LMAITRHGVN RQDTGPLMKC SFEETVDVLM
EAAAHGESDP MKGVSENIML GQLAPAGTGC FDLLLDAEKC KYGMEIPTNI PGLGAAGPTG
MFFGSAPSPM GGISPAMTPW NQGATPAYGA WSPSVGSGMT PGAAGFSPSA ASDASGFSPG
YSPAWSPTPG SPGSPGPSSP YIPSPGGAMS PSYSPTSPAY EPRSPGGYTP QSPSYSPTSP
SYSPTSPSYS PTSPNYSPTS PSYSPTSPSY SPTSPSYSPT SPSYSPTSPS YSPTSPSYSP
TSPSYSPTSP SYSPTSPSYS PTSPSYSPTS PSYSPTSPSY SPTSPSYSPT SPSYSPTSPS
YSPTSPNYSP TSPNYTPTSP SYSPTSPSYS PTSPNYTPTS PNYSPTSPSY SPTSPSYSPT
SPSYSPSSPR YTPQSPTYTP SSPSYSPSSP SYSPTSPKYT PTSPSYSPSS PEYTPTSPKY
SPTSPKYSPT SPKYSPTSPT YSPTTPKYSP TSPTYSPTSP VYTPTSPKYS PTSPTYSPTS
PKYSPTSPTY SPTSPKGSTY SPTSPGYSPT SPTYSLTSPA ISPDDSDEEN ASGGYPYDVP
DYACLSYDTE ILTVEYGFLP IGKIVEERIE CTVYTVDKNG FVYTQPIAQW HNRGEQEVFE
YCLEDGSIIR ATKDHKFMTT DGQMLPIDEI FERGLDLKQV DGLPKGDYKD DDDK
```

#### Cell culture

HEK 293T cells were cultured as a monolayer in DMEM (Thermo Fisher), supplemented with 10% v/v FBS (Thermo Fisher), 100 U $ml^{-1}$ penicillin (Thermo Fisher), and 100 µg $ml^{-1}$ streptomycin (Thermo Fisher). Cells were maintained in an incubator at 37 °C with 5% $CO_2$. See *General procedure for photoproximity labeling in nuclei* below for experiments involving small-molecule treatment.

#### Transfection

Each 10 cm plate of HEK293T cells at 70% confluency were transfected with a plasmid encoding POI-HA-$Cfa^N$-FLAG (5 µg per plate) with Lipofectamine 2000 (12 µl per plate) following the manufacturer's instructions. After 6 h, the media was aspirated and replaced with fresh media. Transfection was performed for 24 or 48 h in an incubator at 37 °C with 5% $CO_2$.

#### General procedure for photoproximity labeling in nuclei

**Protein trans-splicing in isolated nuclei for TMT-10 plex experiments.**

$3\times10^7$ HEK 293T cells transfected with POI-HA-Cfa-N-FLAG were lysed by hypotonic lysis in 3 ml RSB buffer (10 mM tris, 15 mM NaCl, 1.5 mM $MgCl_2$, Roche cOmplete EDTA-free protease inhibitors, pH 7.6) for 10 min on ice. The crude nuclei were isolated by centrifugation at 400 g for 5 min at 4 °C. The nuclei were resuspended in 3 ml RSB buffer, and homogenized with ten strokes of a loose pestle Dounce homogenizer, and pelleted at 400 g for 5 min at 4 °C. The nuclei were resuspended in crosslinking buffer (20 mM HEPES, 1.5 mM $MgCl_2$, 150 mM KCl, Roche cOmplete EDTA-free protease inhibitors, pH 7.6) and centrifuged at 400 g for 5 min at 4 °C. Finally, the nuclei were resuspended in 300 µl of crosslinking buffer per 1 $\times10^7$ cells. To the isolated nuclei was added $Cfa^C$-Ir in crosslinking buffer (0.5 $\mu$M final concentration). The nuclei were incubated at 37 °C for 1 h.

The nuclei were isolated by centrifugation at 400 g for 5 min at 4 °C and washed twice with crosslinking buffer (500 µl) to remove excess peptide. The pellets were then resuspended in 3 mL crosslinking buffer containing diazirine-biotin conjugate (200 $\mu$M) and irradiated with blue light for 45s in the Penn PhD Photoreactor M2 at 100% light intensity at 4 °C. The nuclei were re-isolated by centrifugation at 400 g for 5 min at 4 °C and washed once with crosslinking buffer to remove excess biotin-diazirine. The washed pellets were then resuspended in 2 mL LB3 buffer (10 mM tris, 100 mM NaCl, 1 mM EDTA, 0.5 mM EGTA, 0.1% sodium deoxycholate, 0.5% sodium lauroyl sarcosinate, pH 7.5) and sonicated using a Branson probe tip sonicator (12 cycles at 25% amplitude, 15 seconds on 15 seconds off on ice).

The lysed nuclei were then clarified through centrifugation at 15,000 g for 20 mins at 4 °C and the protein concentration of the supernatant was determined by BCA assay. Protein concentration was normalized across all experimental replicates and diluted to 1 mg/mL with binding buffer (25 mM tris, 150 mM NaCl, 0.25% v/v NP-40, pH 7.5). 1 mL of each sample was then incubated with 125 $\mu$L of pre-washed magnetic Sepharose streptavidin beads (https://www.cytivalifesciences.com, No: 28985738) for 2 h at rt with end-over-end rotation. The beads were subsequently washed twice with 1% w/v SDS in PBS, twice with 1 M NaCl in PBS, and 10% EtOH in PBS x 3.

**Experiments with ligand treatment (Figure 4)**

Following transfection, plated cells expressing POI-HA-Cfa-N-FLAG were treated with the small molecule ligand for the specified amount of time (see table below). Following this, cells were scraped and pelleted as described previously. All buffers used in cell processing prior to irradiation (RSB, crosslinking) contained the ligand. Following irradiation, samples were treated as normal.

| Transfection | Ligand | Concentration | Time |
|---|---|---|---|
| H2A-HA-Cfa-N-FLAG | JQ-1 | 5 μM | 3 h |
| H2A-HA-Cfa-N-FLAG | Pinometostat | 2.5 μM | 24 h |
| RPB1-HA-Cfa-N-FLAG | AT7519 | 2 μM | 2 h |

**For western blot analysis**

At this point proteins can be eluted from beads for western blotting by addition of 25 μL of 4x Laemmli sample buffer containing 20 mM DTT and 25 mM biotin and heated to 95°C for 10 min.

**For proteomic analysis**

Following streptavidin-enrichment, the beads were resuspended in PBS (300 μL) and transferred to a new 1.5 mL Lo-bind tube. The supernatant was removed and the beads were washed with 3 x PBS (0.5 mL) and 3 x ammonium bircarbonate (100 mM).

The beads were re-suspended in 500 μL 3 M urea in PBS and 25 μL of 200 mM DTT in 25 mM $NH_4HCO_3$ was added. The beads were incubated at 55°C for 30 min. Subsequently, 30 μL 500 mM iodoacetamide in 25 mM $NH_4HCO_3$ was added and incubated for 30 min at room temperature in the dark. The supernatant was removed and the beads washed with 3 x 0.5 mL DPBS and 6 x 0.5 mL triethyl ammonium bicarbonate (TEAB, 50 mM). The beads were resuspended in 0.5 mL TEAB (50 mM) and transferred to a new protein LoBind tube. The beads were resuspended in 40 μL TEAB (50 mM), 1.2 μL trypsin (1 mg/mL in 50 mM acetic acid) was added and the beads incubated overnight with end-over-end rotation at 37 °C. After 16 hours, an additional 0.8 μL trypsin was added and the beads were incubated for an additional 1 hour at 37 °C. Meanwhile, TMT 10-plex label reagents (0.8 mg) (Thermo) were equilibrated to room temperature and diluted with 41 μL of anhydrous acetonitrile (Optima grade; 5 min with vortexing) and centrifuged. 40 μL of each TMT reagent was added to the appropriate sample. The reaction was incubated for 2 hours at room temperature. The samples were quenched with 8 μL of 5% hydroxylamine and incubated for 15 minutes. The samples were pooled in a new Protein LoBind tube and quenched with TFA (16 µL, Optima). The samples were dried in a SpeedVac before processing by the Princeton Proteomics Core Facility.

Mass spectra were obtained using an Orbitrap Fusion Lumos mass spectrometer at Princeton Proteomics Facility and analysed using MaxQuant. TMT labeled peptides were dried down in a SpeedVac, re-dissolved in 300 µl of 0.1% TFA in water and fractionated into 8 fractions using the Pierce™ High pH Reversed-Phase Peptide Fractionation Kit (#84868).

Fractions 1, 4, and 7 were combined as sample 1. Fractions 2 and 6 were combined as sample 2. Fractions 3, 5, and 8 were combined as sample 3. Three combined samples were dried completely in a SpeedVac and resuspended in 20 µl 5% acetonitrile/water (0.1% formic acid (pH = 3)). 2µl (~ 360 ng protein) was injected per run using an Easy-nLC 1200 UPLC system. Samples were loaded directly onto a 45cm long x 75µm inner diameter nano capillary column packed with 1.9 µm C18-AQ resin (Dr. Maisch, Germany) mated to a metal emitter in-line with an Orbitrap Fusion Lumos (Thermo Scientific, USA). The column temperature was set at 45 °C and a two-hour gradient method running at a flow rate of 300 $nlmin^{-1}$ was used. The mass spectrometer was operated in data dependent mode with synchronous precursor selection (SPS) - MS3 method[5] with 120,000 resolution MS1 scan (positive mode, profile data type, intensity threshold $5.0e^3$ and mass range of 375-1600 m/z) in the Orbitrap followed by CID fragmentation in the ion trap with 35% collision energy for MS2 and HCD fragmentation in the Orbitrap (50,000 resolution) with 55% collision energy for MS3. The MS3 scan range was set to 100-500 with injection time of 120 ms. A dynamic exclusion list was invoked to exclude previously sequenced peptides for 60 s and a maximum cycle time of 2.5 s was used. Peptides were isolated for fragmentation using the quadrupole (0.7 m/z isolation window). The ion-trap was operated in Rapid mode.

MS/MS/MS data was searched against the Uniprot human protein database containing common contaminants. Each TMT 10-plex experiment was loaded as three fractions and database search criteria were applied as follows: variable modifications set to methionine oxidation and *N*-terminal acetylation and deamidation (NQ), and fixed modifications set to cysteine carbamidomethylation, with a maximum of 5 modifications per peptide. Specific tryptic digestion (trypsin/P) was selected with a maximum of 2 missed cleavages. Match between runs was selected for fractions obtained from the same TMT 10-plex experiment. The maximum peptide mass was set to 6000 Da. The label minimum ration count was set to 2 and quantified using both unique and razor peptides. FTMS MS/MS match tolerance was set to 0.05 Da, and ITMS MS/MS match tolerance was set to 0.6 Da.  All other settings were left as default.

The proteinGroups.txt file was subsequently imported into Perseus (https://maxquant.net/perseus/ ) The data were subsequently filtered based upon the following criteria, 'only identified by site', 'reverse', and 'potential contaminant'. The resulting data was Log2 transformed and median normalization was performed. FDR-corrected p values were determined by a 2-sample T-test following the Benjamini-Hochberg procedure. The data were visualized by plotting as a volcano plot.

#### Procedure for mononucleosome IP (Figure 3e)

The following protocol was adapted from Khan, K. A. *et al. Front. Cell Dev. Biol.* **2020**, *8*, #331, with minor modifications, as described below.

Two 10 cm plates of HEK293T cells transfected with H2A-E92K-HA-Cfa$^{N}$-FLAG (approx. 30 million cells) and two 10 cm plates of HEK203T cells transfected with H2A-HA-Cfa$^{N}$-FLAG (approx. 30 million cells) were lysed in 1 ml hypotonic lysis buffer (10 mM tris, 15 mM NaCl, 1.5 mM $MgCl_2$, Roche cOmplete EDTA-free protease inhibitors, 1 mM DTT, 5 mM sodium butyrate, pH 7.6) for 10 min on ice, and the nuclei were pelleted at 400 g for 5 min at 4 °C.

The nuclei were then resuspended in RSB + 1 mM DTT + 0.2% v/v triton + 5 mM Na butyrate (500 µL per condition) and incubated on ice for 5 min. Nuclei were then pelleted at 400 g for 5 min at 4 °C. The nuclei were washed once more with with RSB + 1 mM DTT + 5 mM Na butyrate and centrifuged at 600 g for 5 min at 4 °C.

The nuclei were then resuspended in 500 µl MNase digestion buffer (10 mM tris, 60 mM KCl, 15 mM NaCl, 2 mM $CaCl_2$, pH 7.5) and were incubated at 37 °C for 10 min. MNase (5µL, NEB) was then added to each condition for 10 min at 37 °C (*N.B.* digestion time varies by enzyme batch and must be determined for each experiment. 2 µl of the digest was removed every two minutes and quenched by addition of 20 mM EGTA. These aliquots were run on a 1.2% agarose gel and the digestion efficiency visualized with ethidium bromide staining). Once digestion to mononucleosomes was complete, the reaction was quenched with the addition of 20 mM EGTA on ice for 5 min. The sample was spun at 1.3 kg for 5 min at 4 °C and the supernatant was collected (fraction S1). 500 µL of buffer TE + 5 mM Na butyrate (10 mM tris, 1 mM EDTA, pH 8.0) was added to the pellet and the sample was rotated end-over-end at 4 °C for 30 min. The sample was spun at 13,000 g for 5 min at 4 °C and the supernatant was collected (fraction S2).

To 475 µL of S1 was added 475 µL of 2× buffer E (30 mM HEPES, 225 mM NaCl, 3 mM $MgCl_2$, 0.4% triton X-100, 20% v/v glycerol, pH 7.5) with constant vortexing dropwise over one minute. To 475 µl of S2 was added 237 µL of 3× buffer D (60 mM HEPES, 450 mM NaCl, 4.5 mM $MgCl_2$, 0.6 mM EGTA, 0.6% triton X-100, 30% v/v glycerol, pH 7.5) with constant vortexing dropwise over one minute. The samples were spun at 13,000 g for 5 min at 4 °C. The two fractions were combined in a 5 ml Lo-bind Eppendorf tube and 30 µl of magnetic FLAG beads (Sigma M2 anti-FLAG magnetic beads; pre-washed with 1× buffer D) were added per condition. The FLAG-IP was performed overnight with end-over-end rotation at 4 °C.

The beads were washed sequentially with 1× buffer D and 1× buffer D + 0.5% v/v triton X-100 for 2 min each with rotation. 40 µL 1× SDS loading buffer was added and the beads were boiled for 10 min. Samples were run on a 15% tris gel for western blotting with appropriate antibodies.

#### Western blotting

SDS-PAGE gels were transferred to PVDF membranes and blocked with 3% w/v bovine serum albumin in TBS-T (25 mM tris, 150 mM NaCl, 0.1% v/v tween-20, pH 7.7) for 30 min at room temperature. Membranes were incubated with primary antibodies at the stated dilutions (**see Methods**) overnight with rotation at 4 °C. After 3 x 5 min washes with TBS-T, secondary antibodies were applied for one hour at room temperature (**See Methods**), before imaging on a Li-

Cor Odyssey imager (LI-COR; IRDye secondaries) or an ImageQuant LAS-4000 (GE Healthcare; HRP secondaries).

#### Synthetic procedures

**3-(4'-Methyl-[2,2'-bipyridin]-4-yl)propanoic acid**

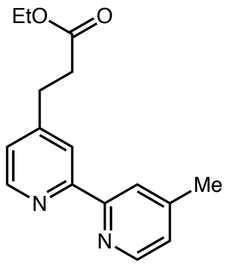

3-(4'-Methyl-[2,2']bipyridinyl-4-yl)-propionic acid ethyl ester**.** 4,4'-Dimethyl-2,2'-bipyridyl (2.5 g, 13.5 mmol) was dissolved in dry THF (20 mL) under a nitrogen atmosphere in a flame-dried flask. The solution was cooled to -78 °C, and a solution of lithium diisopropylamide (14.8 mmol, 1.1 equiv) was added. The reaction mixture was allowed to warm to room temperature for 1.5 hours. This solution was cannulated into a solution of ethyl 2-bromoacetate (2.3 ml, 20 mmol) in dry THF (15 ml) at -78 °C under $N_2$. The reaction mixture was allowed to reach room temperature slowly overnight and quenched by addition of sat. sodium bicarbonate solution. Work-up using ethyl acetate followed by drying over $Na_2SO_4$ and concentration under reduced pressure provided the crude product. The crude residue was purified by column chromatography (Silica gel; DCM:MeOH:$NH_4OH$ 95:5:0.5) to provide the desired product in 69% yield. Spectral data was consistent with literature reports.

**3-(4'-methyl-[2,2'-bipyridin]-4-yl)propanoic acid**

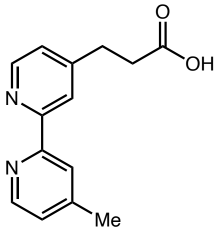

The bipyridinyl ethyl ester was taken up in 1:1 THF:water before the addition of LiOH (2 equiv.). The reaction mixture was stirred at room temperature for 16 h (completion by TLC) before being quenched through the addition of $NH_4Cl$ (until pH 5-6). The mixture the extracted with EtOAc, dried over $Na_2SO_4$ and concentrated under reduced pressure to provide the desired product as an off-white powder (63% yield). Spectral data was consistent with literature reports.

**Ir-$CO_2$H**

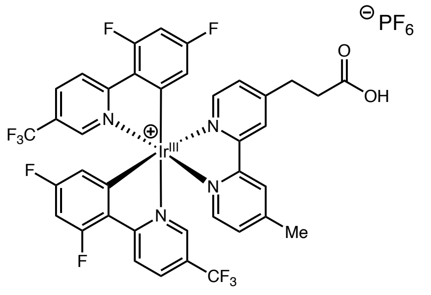

To a round bottomed flask charged with 3-(4'-methyl-[2,2'-bipyridin]-4-yl)propanoic acid and Ir[dF($CF_3$)ppy]$MeCN_2$ $PF_6$ was added DCM/EtOH (4:1) and the reaction mixture was stirred at 30 °C for 16 hours. The resulting solution was concentrated under reduced pressure to provide a yellow solid. The crude product was purified by flash column chromatography (silica gel, 0-10% MeOH/DCM) to provide the desired acid bearing Ir-catalyst (**34**) (55% yield).

**$^1$H NMR (500 MHz, *d*$_6$-DMSO):** δ 12.31 (br, 1H), 8.84 (br, 2H), 8.46 (m, 4H), 7.83 (dd, 2H, *J* = 10.2, 5.7 Hz), 7.67 (s, 1H), 7.65 (dd, 1H, *J* = 5.8, 1.7 Hz), 7.59 (m, 1H), 7.50 (s, 1H), 7.08 (ddd, 2H, *J* = 12.2, 9.3, 2.3 Hz), 5.78 (td, 2H, *J* = 8.1, 2.4 Hz), 3.07 (t, 2H, *J* = 7.5 Hz), 2.75 (t, 2H, *J* = 7.5 Hz), 2.58 (s, 3H)

**$^{13}$C NMR (126 MHz, *d*$_6$-DMSO):** δ 173.58, 167.24, 165.34 (d, *J* = 13.0 Hz), 163.27 (dd, *J* = 13.0, 6.0 Hz), 161.16 (d, *J* = 13.4 Hz), 156.01, 155.80 (t, *J* = 6.4 Hz), 155.61, 155.42, 153.06, 145.74 (d, *J* = 54.4 Hz), 138.04, 130.13, 129.40, 126.91, 126.44, 125.56, 125.01 (dd, *J* = 36.2, 2.7 Hz),

124.15 (d, $J$ = 20.4 Hz), 122.38 (dd, $J$ = 272.2, 6.8 Hz), 114.53 (t, $J$ = 15.7 Hz), 100.07 (t, $J$ = 27.0 Hz), 33.5, 30.2, 21.4.

**$^{19}$F NMR (376 MHz, $d_6$-DMSO):** δ -61.4 (s, 3F), -61.6 (s, 3F), -70.1 (d, 6F, $J$ = 711.3 Hz), -103.3 (ddt, 2F, $J$ = 26.6, 12.1, 8.9 Hz), -106.8 (t, 2F, $J$ = 12.3 Hz)

**HRMS (ESI-TOF)** m/z calcd. for $[M]^+$ = 951.1369, found 951.1332

**Ir-DBCO**

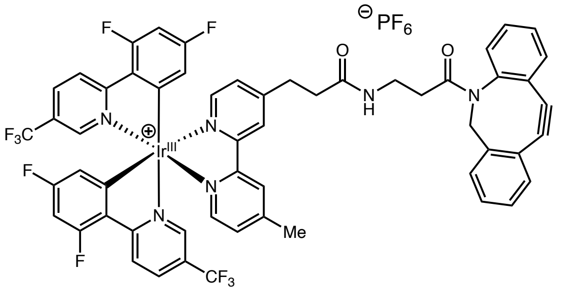

In the dark, a vial was charged with Ir-$CO_2$H (1 equiv) and PyBop (2 equiv.). The vial was purged by through evacuation under vacuum followed by backfilling with nitrogen. The process was repeated three times and the vial was left under nitrogen. To this was added DMF (1 mL) and EtN*i*$Pr_2$ (3 equiv.) and the reaction mixture was sparged with nitrogen for 30 mins. To a separate vial charged with DBCO amine was added DMF (1 mL) and was sparged for 30 mins. After sparging, the amine solution was added to the acid under nitrogen and the solution was stirred for 16 h. *Important: the reaction must be run in the complete exclusion of light.* Upon completion of the reaction, the solution was diluted with EtOAc and washed with NaHCO3 (x2), 0.1 M HCl (x2) and brine (x2). The organics were then dried over sodium sulfate and concentrated at reduced pressure to give a yellow solid. The crude product was purified by trituration with cold $Et_2O$ to provide the desired catalyst (80% yield).

**$^1$H NMR (500 MHz, $d_6$-DMSO):** δ 8.80 (dd, $J$ = 12.2, 7.5 Hz, 2H), 8.44 (p, $J$ = 10.2 Hz, 4H), 7.87 – 7.80 (m, 2H), 7.79 (d, $J$ = 7.2 Hz, 1H), 7.65 (dd, $J$ = 15.9, 6.7 Hz, 2H), 7.61 – 7.57 (m, 2H), 7.55 – 7.50 (m, 2H), 7.45 (h, $J$ = 4.3 Hz, 2H), 7.38 (t, $J$ = 7.5 Hz, 1H), 7.31 (t, $J$ = 7.6 Hz, 1H), 7.24 (t, $J$ = 8.0 Hz, 1H), 7.10 – 7.00 (m, 2H), 5.83 – 5.71 (m, 2H), 5.04 (dd, $J$ = 14.0, 3.3 Hz, 1H), 3.63

(dd, $J$ = 14.0, 8.4 Hz, 1H), 3.14 – 2.85 (m, 5H), 2.57 (s, 3H), 2.47 – 2.35 (m, 2H), 1.93 – 1.81 (m, 1H).

**$^{13}$C NMR (126 MHz, $d_6$-DMSO):** δ 170.2 (d, $J$ = 8.5 Hz), 166.8, 164.9 (d, $J$ = 12.8 Hz), 162.8 (dd, $J$ = 13.4, 5.8 Hz), 160.7 (d, $J$ = 13.5 Hz), 156.1 (d, $J$ = 3.7 Hz), 155.4 (d, $J$ = 6.6 Hz), 155.21 – 155.0 (m), 152.7, 151.4, 150.1 (d, $J$ = 31.3 Hz), 148.4, 145.3 (d, $J$ = 43.7 Hz), 137.6, 132.4 (d, $J$ = 2.5 Hz), 129.6 (d, $J$ = 19.9 Hz), 128.9 (d, $J$ = 11.8 Hz), 128.2 (d, $J$ = 20.2 Hz), 127.7, 126.8 (d, $J$ = 3.7 Hz), 126.0, 125.2 (d, $J$ = 12.1 Hz), 124.6 (q, $J$ = 34.6 Hz), 123.7 (d, $J$ = 20.4 Hz), 123.0 (d, $J$ = 3.8 Hz), 122.5 (d, $J$ = 5.1 Hz), 121.5, 120.8 (d, $J$ = 3.0 Hz), 114.3 (d, $J$ = 3.3 Hz), 114.1 (t, $J$ = 15.1 Hz), 108.0 (d, $J$ = 7.0 Hz), 99.6 (t, $J$ = 26.9 Hz), 54.9 (d, $J$ = 11.6 Hz), 45.9 (d, $J$ = 4.1 Hz), 35.1 (d, $J$ = 3.5 Hz), 34.7 (d, $J$ = 4.1 Hz), 34.2 (d, $J$ = 3.8 Hz), 30.4 (d, $J$ = 5.8 Hz), 25.9 (d, $J$ = 7.6 Hz), 21.0.

**$^{19}$F NMR (376 MHz, $d_6$-DMSO):** δ -61.53 – -61.63 (m), -61.68 (d, $J$ = 3.8 Hz), -69.24, -71.13, -103.31 (ddd, $J$ = 13.0, 9.7, 7.3 Hz), -106.77 (q, $J$ = 11.3 Hz).

**HRMS (ESI-TOF)** m/z calcd. for $[M]^+$ 1209.2520, found 1209.2531

**Ir-NHS**

To 100 μL of a 10 mM DMSO solution of Ir-DBCO was added 10 μL of a 100 mM stock solution (DMSO) of azido-PEG16-NHS ester (Broadpharm). The mixture was rotated end over end for 2 hours. Analysis by HRMS showed complete conversion to the SPAAC product, the Ir-NHS ester was used immediately for Ir-intein solid phase synthesis.

**HRMS (ESI-TOF)** m/z calcd. for $[M]^+$ 2125.72657, found 2125.72789.

### Supplementary Tables

#### Supplementary Table 1-14. Proteomics hits

Supp. Tables 1-14 are provided as XL files and contain all protein hits for each μMap experiment.

### Uncropped western blots for data presented in Figures 2&3

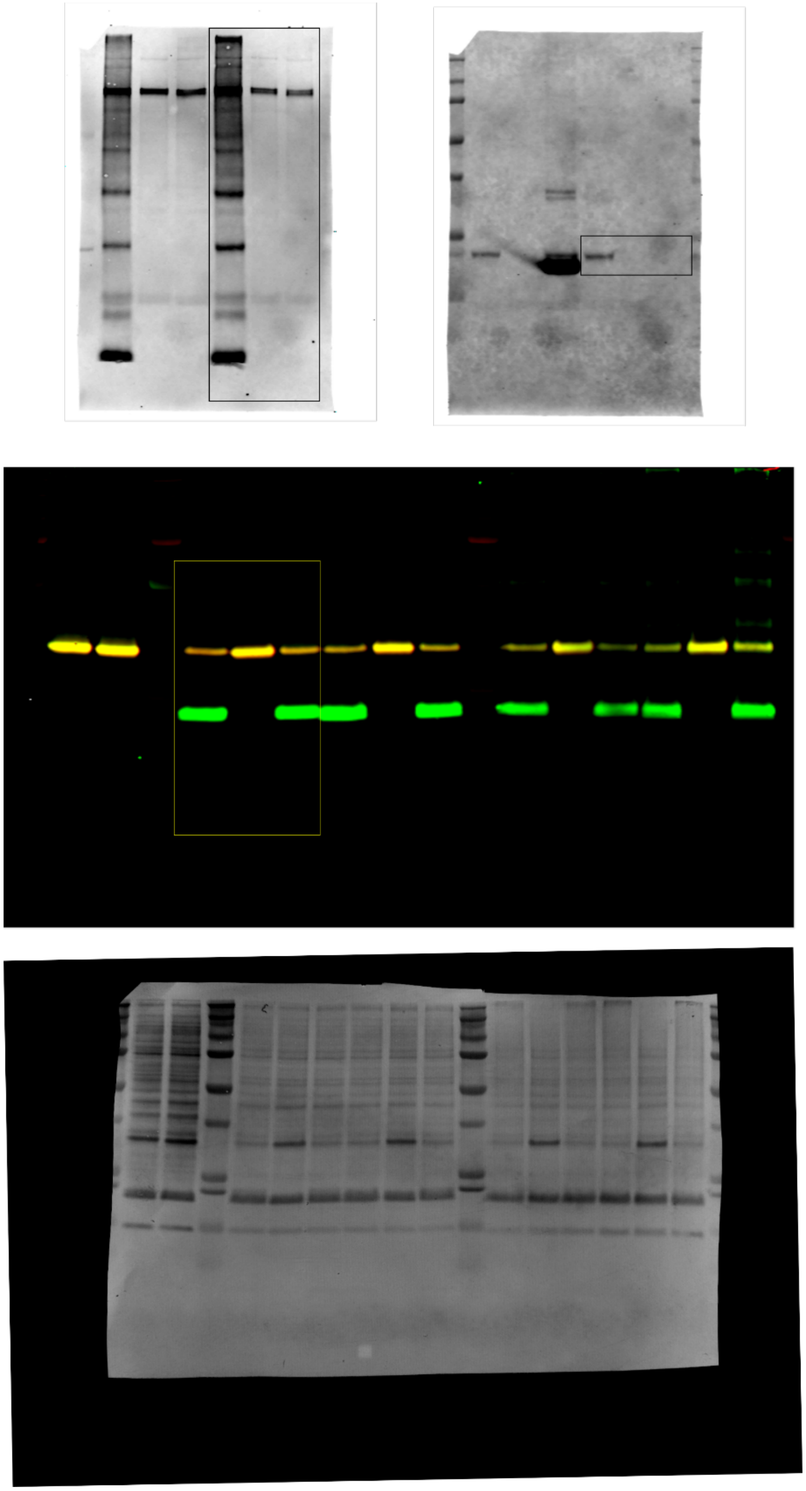

Uncropped western blots for data presented in Figure 2.

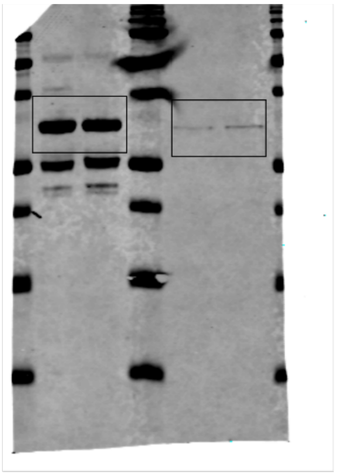

anti-FLAG

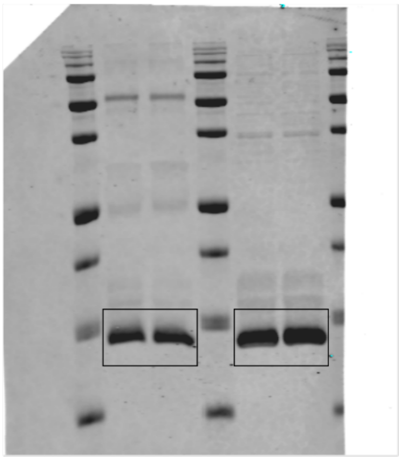

anti-H4

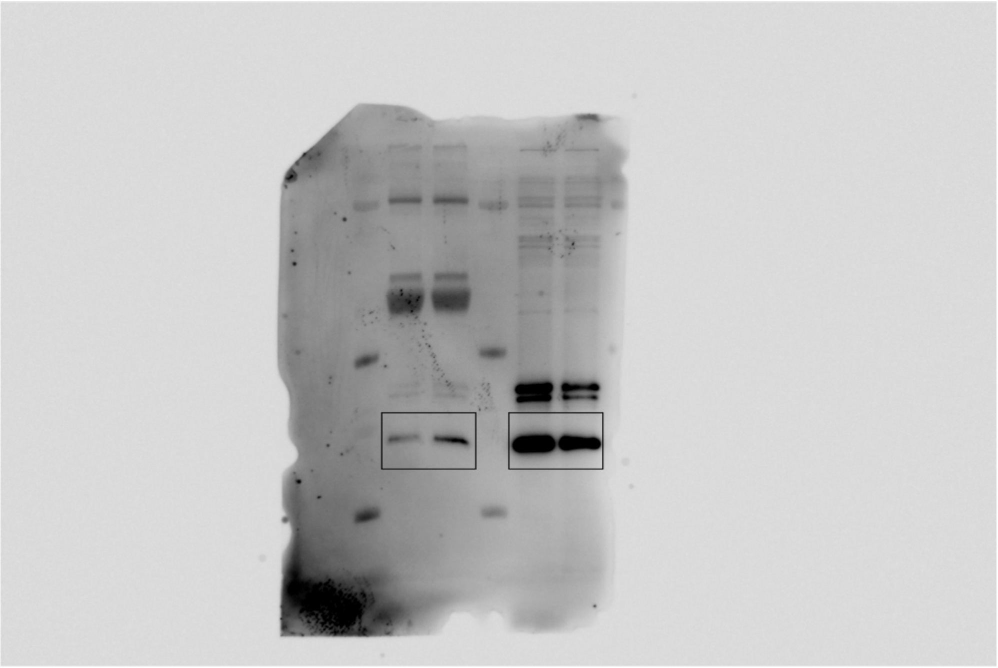

anti-Acetyl (pan)

Uncropped western blots for data presented in Figure 3.
